# Using EEG Microstates to Examine Post-Encoding Quiet Rest and Subsequent Word-Pair Memory

**DOI:** 10.1101/2020.05.08.085027

**Authors:** Craig Poskanzer, Dan Denis, Ashley Herrick, Robert Stickgold

## Abstract

Evidence suggests that the brain preferentially consolidates memories during “offline” periods, in which an individual is not performing a task and their attention is otherwise undirected, including spans of quiet, resting wakefulness. Moreover, research has demonstrated that factors such as the initial encoding strength of information influence which memories receive the greatest benefit. Recent studies have begun to investigate these periods of post-learning quiet rest using EEG microstate analysis to observe the electrical dynamics of the brain during these stretches of memory consolidation, specifically finding an increase in the amount of the canonical microstate D during a post-encoding rest period. Here, we implement an exploratory analysis to probe the activity of EEG microstates during a post-encoding session of quiet rest in order to scrutinize the impact of learning on microstate dynamics and to further understand the role these microstates play in the consolidation of memories. We examined 54 subjects (41 female) as they completed a word-pair memory task designed to use repetition to vary the initial encoding strength of the word-pair memories. In this study, we were able to replicate previous research in which there was a significant increase (p < .05) in the amount of microstate D (often associated with the dorsal attention network) during post-encoding rest. This change was accompanied by a significant decrease (p < .05) in the amount of microstate C (often associated with the default mode network). We also found preliminary evidence indicating a positive relationship between the amount of microstate D and improved memory for weakly encoded memories, which merits further exploration.

## 1. Introduction

Following initial encoding, memories undergo a consolidation process to strengthen and transform them into enduring, long-term, memory traces. “Offline” periods are beneficial to this process; specifically, a large body of research shows that sleep is critical for memory consolidation [Stickgold 2005, Walker and Stickgold 2004, Karni et al. 1994]. A growing collection of studies indicates that spans of resting wakefulness also promote the consolidation of memories [for a review see Wamsley, 2019]. These periods are task-negative intervals (periods without a specific task and with reduced attentional demand, as opposed to task-positive intervals) during which new encoding is limited and the brain is otherwise occupied by mind-wandering [Wamsely, 2019]. Quiet rest has been shown to protect the fine-grained details of labile memories in a visual recognition task [Craig and Dewar 2018], as well to improve recollection of auditory stimuli, as compared to spending an equivalent time performing a distractor task [Brokaw et al. 2016]. Moreover, in studies of mice, researchers have demonstrated that wakeful hippocampal sharp-wave ripples contribute to the accuracy of stored spatial memories [Tang et al. 2017]. Importantly, this rest-related benefit to memory is also seen for hard-to-rehearse information, suggesting that quiet-rest consolidation is not simply due to additional time to internally rehearse learned information [Wamsely, 2019; Humiston et al., 2018; and Dewar et al. 2014]. In support of this conclusion, both Brokaw et al. [2016] and Gregory et al. [2014] noted that spontaneously thinking about recently encoded information during rest does not correlate with improved memory, emphasizing that the rest-related benefits to memory consolidation are not dependent on conscious thought [Brokaw et al., 2016; Gregory et al. 2014]. This body of work illustrates that although quiet rest provides a benefit to the consolidation of memories, the mechanisms facilitating this interaction are not entirely clear. To this end, post-encoding quiet rest periods pose a unique opportunity to provide insight into how the brain processes memories while offline.

Recent research has suggested that one contributing factor to the modulation of memory consolidation across sleep is the initial encoding strength of the memory [sleep benefits weakly encoded memories: Drosopoulos et al., 2007; Schapiro et al., 2017; Djonlagic et al., 2009; Denis et al., 2019; Denis et al., 2020; sleep benefits strongly encoded memories: Tucker and Fishbein, 2008; Schoch et al., 2017; and Wislowska et al., 2017; sleep benefits encoded memories on an inverse U shaped curve: Stickgold, 2009]. In this vein, there are differing factions of evidence— some suggesting sleep benefits weakly encoded memories and others suggesting a benefit for stronger encodings. Studies finding a benefit for weakly encoded memories often used the frequency of stimulus presentation as a measure of encoding strength [Drosopoulos et al., 2007; Schapiro et al., 2017; Denis et al., 2019]; in contrast, studies showing a benefit for strongly encoded information manipulated encoding strength by adding an initial recall test as an opportunity for additional interaction with the task stimuli (in this case, emotional images instead of word-pairs) [Schoch et al., 2017]. Considering whether quiet rest consolidation acts preferentially over similar salience cues as sleep-dependent consolidation is an important question that has recently begun to be explored. In a 2018 paper, *Schapiro et al*. found that weakly encoded memories for object features were replayed in the hippocampus more often than strongly encoded information over the course of a 6.5-minute period of quiet rest [Schapiro et al., 2018]. For this reason, continuing to investigate how the brain selects memories for consolidation during quiet rest is an important next-step in understanding the types of information that are benefitted by offline consolidation periods.

In order to study how memories are prioritized for consolidation over quiet rest, it is important to choose an analysis technique that allows for in-depth scrutiny of the fine-grained temporal dynamics of underlying brain activity. EEG microstates are an ideal tool for examining resting-state brain activity. Microstates are quasi-stable topographical maps of EEG electric potential [Lehmann, 1987]. Across subjects and studies, when plotting the maps of voltage over the scalp, four canonical patterns emerge which are typically referred to as microstates A through D [Michel and Koenig 2018]. These patterns of electric potential remain stable for periods of 80-120 milliseconds before rapidly switching directly to another microstate, rather than gradually morphing between patterns over the course of time [Michel and Koenig 2018; Van De Ville, Britz, and Michel 2010]. Microstates are believed to be measures of global network activity, and have been associated with spikes in activation of various functional networks [Britz et al, 2010]. In their 2010 study, Britz and colleagues demonstrated how each of the canonical microstates mapped onto existing fMRI networks: microstate A aligned with a phonological processing network, microstate B with a visual network, microstate C correlated with task-negative activity and activity in areas that have now been identified with the default mode network (DMN)[Britz, Van De Ville, and Michel 2010, Sestieri et al., 2011; Damoiseaux et al., 2008; Lei et al., 2014; and Xu et al., 2016], and microstate D matched the dorsal attention network, a task-positive system that is active while performing attention-demanding activities [Britz, Van De Ville, and Michel 2010; Corbetta and Shulman, 2002; Mantini et al., 2007; and Damoiseaux et al., 2006]. Further elucidating the relationship between microstate time courses and existing fMRI networks, Van De Ville et al. showed that microstates demonstrate monofractal scale-free dynamics, meaning the smoothed order of microstates switching back and forth is consistent across time scales from 256 ms—the scale of EEG recordings— to 16 seconds— the scale of fMRI recordings. Thus, EEG microstate dynamics are reflected (in fact statistically indistinguishable) in the BOLD signal dynamics seen in fMRI [Van De Ville, Britz, and Michel 2010]. As “atoms of thought” [Michel and Koenig, 2018] microstates offer a high temporal resolution view of cognitive activity during quiet rest. This analysis provides a unique perspective on the underlying network activity that gives rise to the consolidation of memory.

Although there has not been much research on this topic, one recent paper by Murphy et al. [2018] began to explore microstate dynamics in stretches of post-encoding quiet rest [Murphy et al., 2018]. In their study, Murphy and colleagues found that microstate D increased over baseline levels during a seven-minute period of post-encoding rest, an increase that was not seen in subjects who underwent a control non-learning task before quiet rest [Murphy et al., 2018]. Further exploring this relationship, however, they found that this increase in microstate D was negatively correlated with post-nap memory improvement in a virtual maze task, while there were no significant interactions between microstate D and memory performance in subjects who spent the equivalent time awake [Murphy et al., 2018].

In addition to microstate analysis, spectral power has long been a tool to study neural activity during memory tasks. In particular, alpha band frequency (8-13Hz) is believed to be related to memory [Vogt et al., 1998], although results across studies have provided conflicting evidence for the exact relationship between alpha frequency and memory consolidation. In a study by Brokaw et al. [2016], a post-learning quiet rest was associated with a decrease in alpha power; moreover, alpha power during this rest was negatively correlated with later memory performance [Brokaw et al. 2016]. Contrastingly, in a separate study by Murphy et al. [2018], post-encoding rest was associated with increased alpha power in a spatially localized group of electrodes [Murphy et al., 2018]. These asymmetrical results suggest an important area for further exploration in order to understand how spectral power in post-encoding rest relates to memory consolidation.

In this current study, we used microstate analysis to study the network dynamics in a post-encoding rest period. We recorded subjects’ EEG activity during three sessions of eyes-closed quiet rest: one baseline session, one post-encoding, and a final session after a cued recall test of the encoded material. Between rests, the subjects performed a word-pair memory task in which the stimuli were presented for varying lengths of time and with varying amounts of repetition in order to modulate their initial encoding strength. We explored whether there was a shift in resting-state EEG activity after periods of performing this word-pair task and whether these changes were related to subsequent memory performance for word-pairs across encoding strengths.

## 2. Methods

Two separate studies, designed with similar procedures, were performed to identify how encoding strength modulates later consolidation of the memory. The second study, is reported in detail in the Supplementary Results section (see Supplementary Results, Continuous Strength of Encoding Study). Data are available from the corresponding author upon reasonable request.

### 2.1 Participants

A total of 54 subjects (41 female; mean age = 22, SD = 3 years) participated in this study; one subject was excluded as a result of a corrupted data file, additionally, one subject was excluded for having been unable to correctly remember any word pairs from the learning task. All participants were screened to have no history of sleep, neurological, or psychiatric disorders, typical bedtimes no later than 2 a.m., and an average of at least 6 hours of sleep per night. Participants were instructed to maintain a regular sleep schedule for 3 nights leading up to the study and to refrain from consuming any caffeine on the morning of the experiment. Recruitment was through online postings at local universities advertising a learning and memory study. Participants were compensated for their time. Procedures were approved by the institutional review board of Beth Israel Deaconess Medical Center. A portion of this dataset has been previously published by Denis et al. 2020 [Denis et al. 2020, preprint].

### 2.2 Procedure

After providing informed consent, subjects completed a series of questionnaires to determine their sleep quality over the past 3 nights, their sleep habits over the preceding month (Pittsburgh Sleep Quality Index (PSQI)), and their current level of subjective sleepiness/alertness (Stanford Sleepiness Scale (SSS)) [Buysse et al., 1989; Hoddes et al., 1972; Hoddes et al., 1973]. After providing consent and filling out the questionnaires, participants were wired with EEG (detailed below).

Subjects began the experimental protocol with 5-minutes of eyes-closed quiet resting wakefulness (QR1) in which they were not directed to think about anything. Next was an encoding session, detailed below, in which subjects were shown each word-pair a given number of times based on its encoding strength classification. Immediately after seeing the word pairs, subjects had a second, post-encoding, eyes-closed quiet rest (QR2). This was followed by a mind-wandering questionnaire (Amsterdam Resting-State Questionnaire (ARSQ)) and a blank pie-chart that they completed with the percentages of time during the quiet rest that they thought about each of the following categories: the task, the present, the future, the past, or “other,” [Diaz et al., 2013; Gregory et al., 2016]. The results of these instruments will be discussed in a future publication. Following these questionnaires, participants were given an immediate recall test to assess their memory for the word pairs. At the end of the immediate memory test, participants underwent a third eyes-closed, 5-minute quiet rest (QR3). The data collected after QR3 are the subject of a separate analysis reported elsewhere.

### 2.3 Encoding and Recall Task

All subjects were given a word-pair memory task, in which they were shown 180 word pairs, each containing two random nouns (e.g. lemon—mirror) and then tested in an immediate recall session and, later, a delayed recall session by being given the first word of each pair and being asked to type the pair’s second word. The length of the words used to make each pair was controlled by selecting only words between 4 and 7 letters. The word pairs were divided evenly into three distinct encoding groups: word pairs in the “1 PRES” encoding group were only shown once, while “2 PRES” word pairs were shown twice, and “4 PRES” pairs were shown four times. Before beginning the task, the participants were informed that an effective memorization strategy is to visualize a scene involving the two nouns. For each word-pair presentation, a fixation-cross appeared on the screen for 2,000-3,000ms, followed by the given word pair for 2,000ms, and then a blank screen for 1,000ms. The “jitter” in presentation onset times was designed to minimize contamination of the event-related EEG by preparatory responses (not reported in the present report). After the blank screen that followed each word-pair presentation, participants were asked whether they had successfully visualized a scene involving the two words (“yes” or “no”). Subjects were given a minimum 1-minute break after every 70 trials.

During the recall session, participants were shown the first word from each word pair and asked for its paired word. For each word pair, participants were first presented with a fixation-cross for 2,000-3,000ms, then shown the cue word for 2,000ms and asked to think about its pair. After an additional 2,000-2,500ms, the participants were instructed to enter the corresponding word as quickly and accurately as possible. Participants were told there would be no penalties for guesses or mistakes. All recognizable misspellings (e.g. calender instead of calendar) and mistaken pluralizations (e.g. brains instead of brain) were counted as correct answers. The program reminded the participants to respond if they had not commenced typing after 7-seconds, and it automatically advanced to the next trial after 10-seconds. Each word pair was tested only once. The order of the word-pair presentation and testing were both randomized and the word-pairs in each encoding strength condition were randomized for each subject. All tasks were created and executed with custom scripts written using the Psychtoolbox package for MATLAB [Kleiner et al., 2007].

### 2.4 EEG Recording

EEG was recorded using an AURA-LTM64 amplifier and Twin software (Grass Technologies, Warwick, RI). Data was acquired at a sampling rate of 400Hz using 57 electrodes positioned in accordance with the 10-20 system. Additional electrodes were placed on the collarbone (grounding electrode), the forehead (recording reference), to the left and right of the chin (EMG), below the left and above the right eye (EOG), and on the left and right mastoids. Impedances were kept below 25 KOhm.

All EEG and EOG electrodes were referenced to contralateral mastoids, bandpass filtered between 0.3 and 35Hz, and notch filtered at 60Hz. EMG channels were bandpass filtered between 10 and100Hz. Artifact rejection was then performed on the continuous data based on visual inspection; bad channels were spherically interpolated using the spherical splines algorithm in MATLAB’s EEGLAB toolbox [Delorme and Makeig, 2004]. Finally, an independent component analysis was run on each record and components that reflected eye-movements or faulty channels were rejected based on visual inspection. Please see Supplementary Results, Channel Interpolation and Component Rejection for more details.

### 2.5 Microstate Analysis

Microstate analysis was performed using the Microstate EEGLAB toolbox [Poulsen et al., 2018]. The microstates were generated using the EEG data from the Discrete Strength of Encoding (DSOE) study and later applied to the rest data from the Continuous Encoding Strength (CES) protocol in order to keep the microstate topographies consistent when testing for replicability of the results in the DSOE experiment (see Supplementary Results) [Poulsen et al., 2018; Khanna et al., 2014].

To perform the microstate analysis, the DSOE EEG data were first re-referenced to the average across electrodes and after examination, any bad channels were spherically interpolated [Delorme and Makeig, 2004]. Upon inspection of the records, 19 of 162 records (9 from QR1, 5 from QR2, and 5 from QR3) exhibited an abnormal flattening of the global signal and were removed from the microstate analysis (see Supplementary Results, Signal Flattening); thus, we were unable to perform the microstate analysis on all the records across all 54 subjects who participated in the protocol. Data from the DSOE QR1, QR2, and QR3 were concatenated across all subjects and downsampled by selecting time points at the local peaks of global field potential (when the signal-to-noise ratio is at its highest), before feeding the data into a modified K-means clustering algorithm to calculate the optimal number of microstates to use [Poulsen et al., 2018].

Selection of the number of microstates is based on a combination of maximizing the Global Explained Variance (GEV) (a measure of how well the given microstate topographies explain the variance across the data), and minimizing both the Cross Validation Criterion (CV; a measure of unexplained noise) and the total number of microstate prototypes. We used the modified K-means clustering algorithm to determine the number of microstates, using n = 2:8 [Poulsen et al., 2018].

The microstate segmentation analysis yielded a virtual tie between *n* = 5 microstates (CV = 5.15, GEV = .733) and *n* = 6 microstates (CV = 5.14, GEV = .744; Figure 2). We selected *n* = 5 microstates in order to minimize the number of microstates and then back-fit the five topographies onto the quiet rest data. We focused our analysis on measurements of GEV, as in our previous study [Murphy et al., 2018].

**Figure 1:**
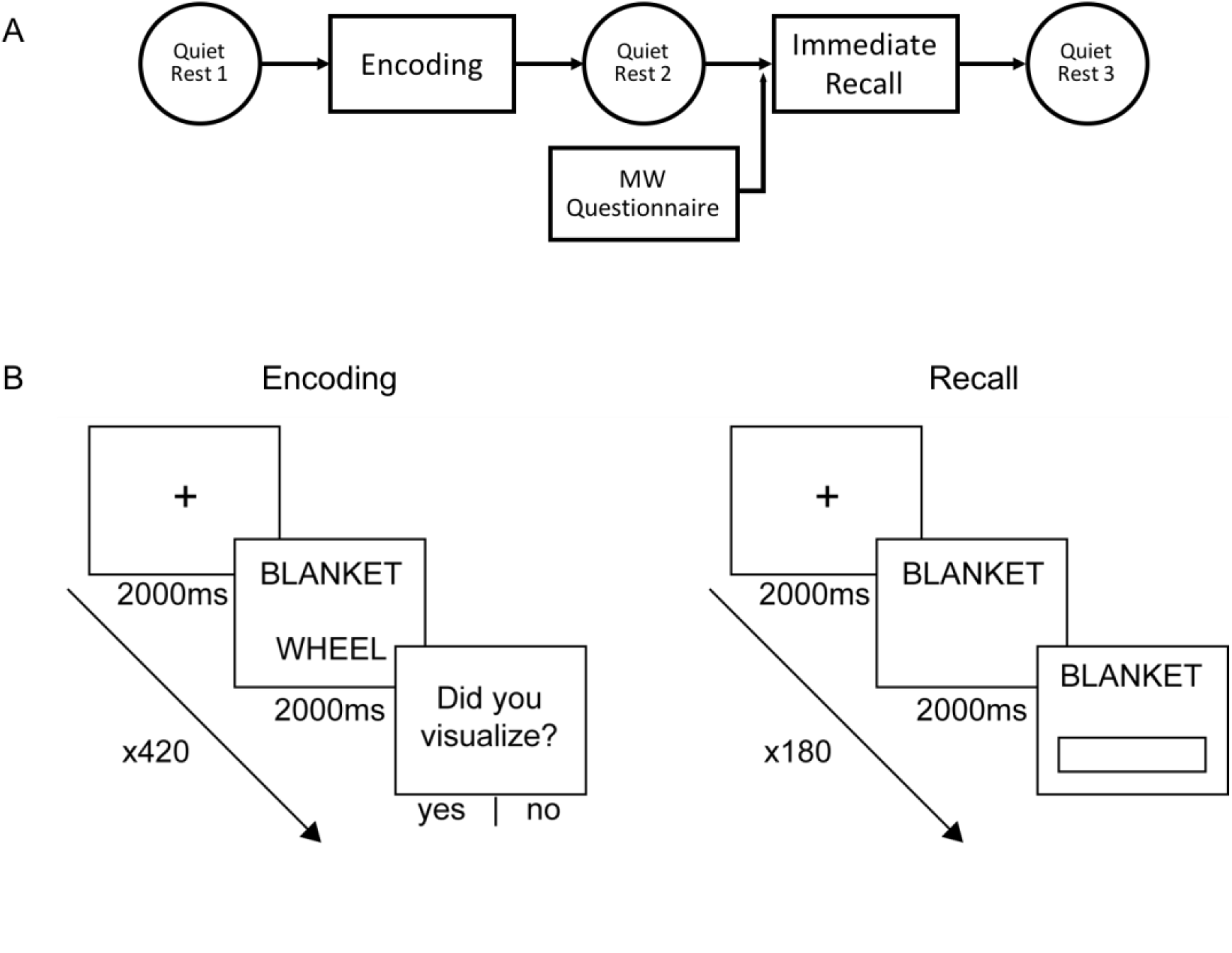
**A:** Protocol for the Discrete Strength of Encoding (DSOE) study. Participants were given 5 minutes of quiet rest before a word-pair memorization task, a second quiet rest (after which they completed a survey detailing their resting thoughts), an immediate cued recall, and a third and final quiet rest. Analyses of the tasks performed after this protocol are analyzed in a separate paper. **B:** Encoding task and recall tasks. In the encoding task, subjects were presented with a pair of random nouns and asked to visualize a scene containing both words in order to improve word-pair memory. In the recall task, subjects were shown one word and asked to type in the word which completes the pair [image adapted from Denis et al. 2020].

**Figure 2:**
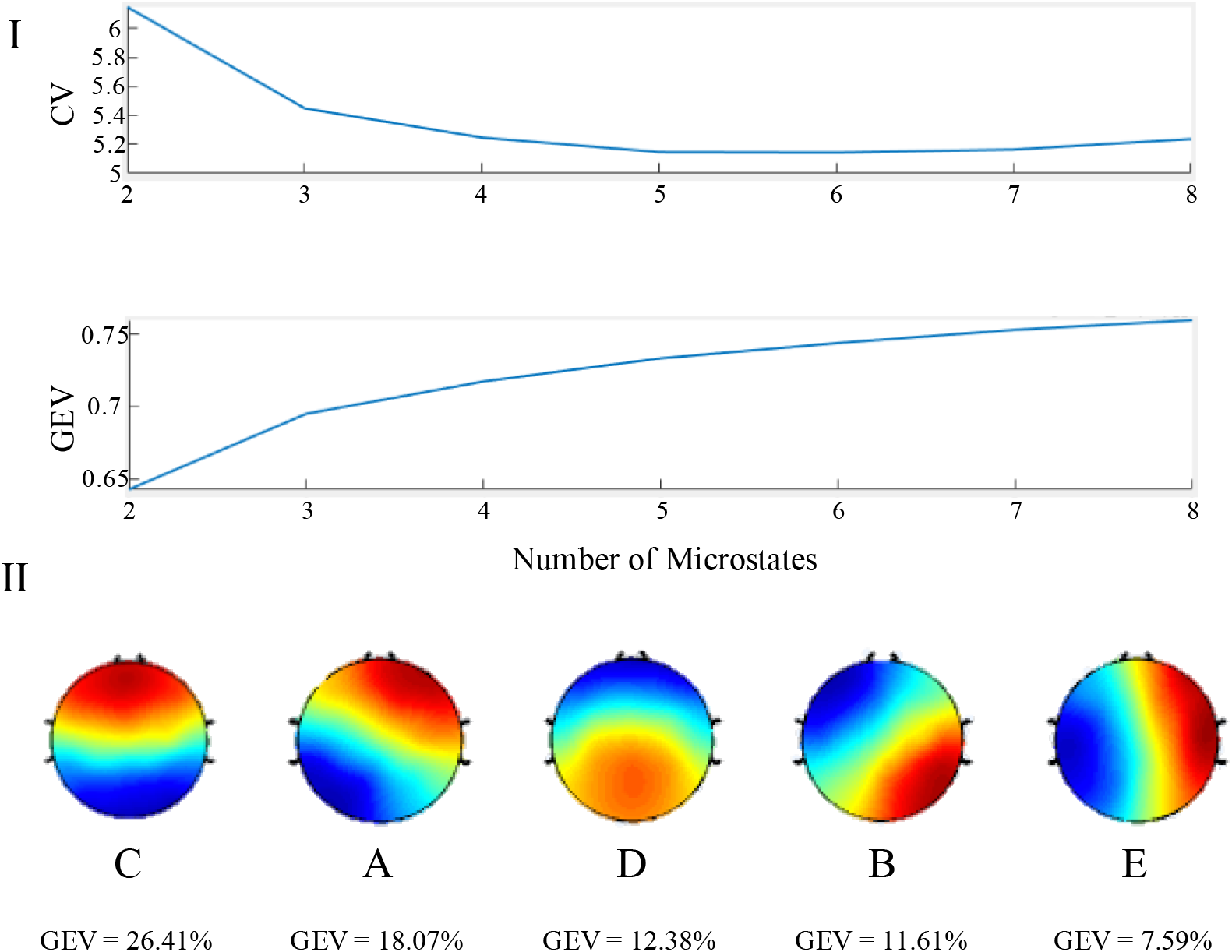
**I:** Selecting an active number of microstates for record segmentation. Top: Graph of the Cross Validation Criterion across active microstates. In order to determine the best fitting number of microstates, the CV should be minimized. Bottom: Graph of the Global Explained Variance across active microstates. Although a larger number of active microstates would explain more global variance, adding additional microstates can lead to over-fitting the data. **II:** Microstate topographies in descending order of GEV. These maps correspond to the 4 canonical microstates in the order of C, A, D, B. The final microstate map (“E”) is a common additional map.

#### 2.5.1 Identifying Microstates

Upon identifying the five active microstates, we compared their topographies to maps of the four canonical microstates (Figure 2). On initial inspection, our set of 5 microstates contained all 4 canonical microstates: topography 1 corresponds to microstate C, topography 2 to microstate A, topography 3 to microstate D, and topography 4 to microstate B. The fifth generated microstate is a common topography among studies that identify more than four active microstates [Custo et al., 2014]. For the sake of consistency, we refer to this fifth map as microstate E.

It is worth noting that there is some variation in the standard topography of microstate D [Michel and Koenig 2018]. Though microstate D is often depicted as having a fronto-central peak [Koenig, 2002], it often appears as an inverse topography with a posterior-central peak [Seitzman, 2017]. In our study, microstate topography 3 matched this inverted map of microstate D, however in *Murphy et al. 2018* the researchers found the standard, fronto-central peak topography of microstate D [Murphy et al. 2018]. In order to remain consistent with previous studies, we refer to both microstate topographies as forms of microstate D.

### 2.6 Spectral Power Analysis

Power spectral density (PSD) was computed at every electrode using the artifact-free data. PSD was derived using Welch’s method (pwelch function in MATLAB [The Mathworks, Natick, MA]) with a 5 second Hamming window and 50% overlap. Spectral power was log-transformed to compensate for the 1/f scaling of the power spectrum. PSD was calculated for the frequency range 1-30Hz. We did not examine spectral power in higher frequency bands (i.e. gamma range) because of low signal-to-noise ratio.

### 2.7 Statistical Analysis

Comparisons between EEG dynamics and behavior in QR1 and QR2 were evaluated using paired sample t-tests. Pearson correlations were used to measure the relationship between EEG and behavioral data. In order to limit the influence of outliers and heteroscedasticity in the data, effects were also analyzed using a robust linear regression model. The false discovery rate (FDR) was used to adjust for multiple comparisons where appropriate. Finally, Meng’s Z-test for comparing correlated coefficients was used to assess the significance of differences in correlation magnitudes [Meng et al., 1992; Spaak, 2020]. All analyses were performed in MATLAB [The Mathworks, Natick, MA] and R [RStudio Team, Boston, MA].

Non-parametric cluster tests were performed on the PSD data to compare the topographic spectral power in baseline (QR1) and post-encoding (QR2) rest as well as between QR2 and memory performance, using the fieldtrip toolbox for MATLAB [Oostenveld et al., 2011]. EEG data from separate experimental conditions are often analyzed by comparing the difference between multiple channels over the course of many time points; cluster-based permutation tests tackle this *multiple comparison problem* by assuming the null hypothesis, that there is no statistical difference between conditions, and repeatedly running t-tests after randomly shuffling the data points between the experimental conditions, creating a distribution of t-values. This t-value distribution demonstrates how likely the original t-value between condition 1 and condition 2 is and thus, if the two conditions are, in fact, statistically distinguishable. In this case, for each time point, EEG data at each channel, at each frequency is repeatedly, randomly relabeled as belonging to either QR1 or QR2 over 10,000 iterations. At each permutation, a between group t-test was executed with a threshold at alpha = 0.05. Channels above the alpha threshold were grouped together and defined as a cluster if the group consisted of 2 or more electrodes. A cluster is deemed significant if its Monte-Carlo Significance probability, the probability of that cluster’s t-statistic appearing over the total permutations, is p < .05.

## 3 Results

### 3.1 Behavior

Immediate recall of word-pairs in the DSOE study increased with initial encoding strength. 77.6% of 4 PRES encoded words were remembered at the initial test as compared to 58.1% for 2 PRES and 31.7% of 1 PRES encoded word pairs. The difference in memory performance across all categories was significant (1 PRES vs. 2 PRES: t(52) = −11.2, p = 1.8 *10^−15^, FDR adjusted p = 2.8 *10^−15^; 2 PRES vs. 4 PRES: t(52) = −7.61, p = 5.3 *10^−10^, FDR adjusted p = 5.3 *10^−10^; 1 PRES vs. 4 PRES: t(52) = −16.0, p = 9.8 *10^−22^, FDR adjusted p = 2.9 *10^−21^).

### 3.2 Microstate fit

For subjects in the DSOE study the five identified microstates explained 79.7% and 77.1% of the global variance for QR1 and QR2, respectively (see Supplementary Results), values consistent with previous studies [Michel and Koenig, 2018]. (The results from QR3 are presented in Supplementary Results).

### 3.3 Microstate Dynamics Across Periods of Rest

In analyzing the DSOE dataset, we found a significant increase in the GEV of Microstate D from baseline rest QR1 to post-encoding rest QR2, increasing from an average of 8.0% to 12.4% (t(38) = 3.2, p = .0027). This increase in microstate D, present in 69% of participants, was accompanied by a significant decrease in the GEV of microstate C in QR2, falling from 24.2% to 19.9% (t(38) = −2.5, p = .016). The decrease in microstate C was present in 67% of participants. Only the increase in microstate D from baseline to post-encoding survived FDR adjustment for multiple comparisons (adjusted p = 0.041; Figure 4). Moreover, we found that microstates C and D GEV were negatively correlated during both QR1 (r = -.45, p = .002, robust p = .004; Supplementary Figure 13, A) and QR2 (r = -.46, p = .001, robust p = .003; Supplementary Figure 13, B), as was the change in microstates C and D from QR1 (r = -.43, p = .006, robust p = .009; Supplementary Figure 13, C).

**Figure 3:**
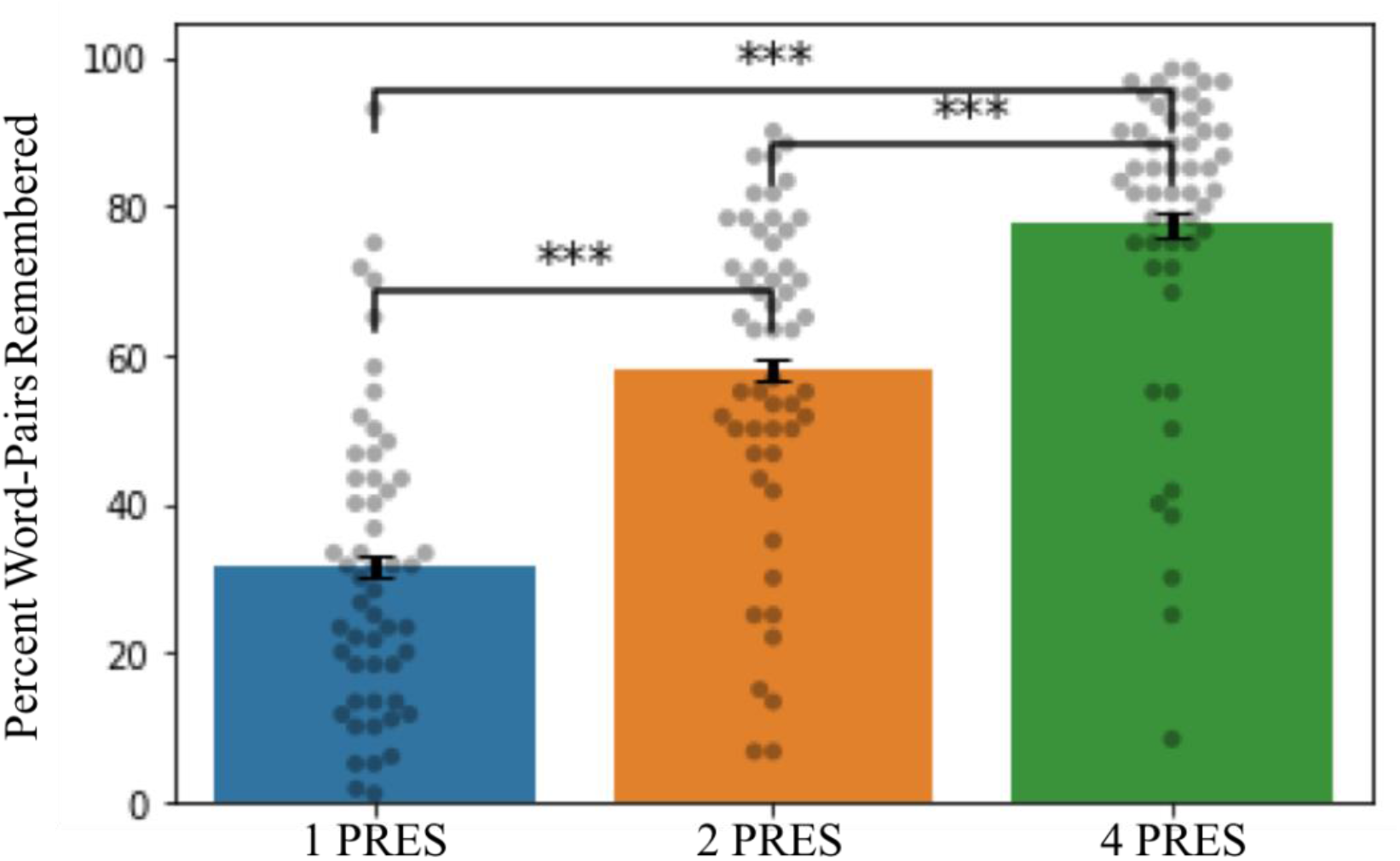
Memory performance in the DSOE study (dots represent individual subject values; error bars display the within subject standard error; ***p < .0001).

**Figure 4:**
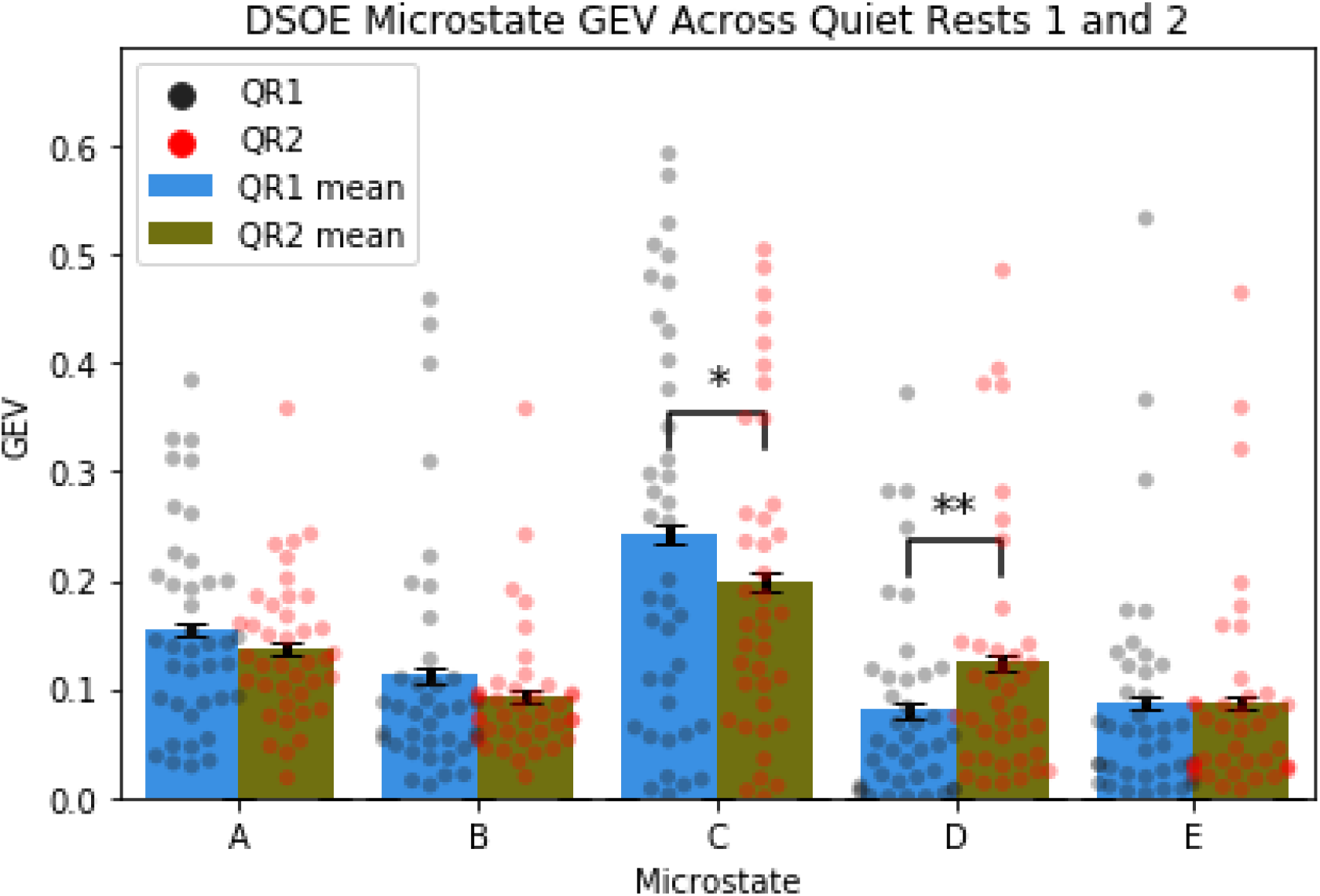
Measuring GEV across rest periods (colored dots represent individual subject values across conditions; error bars display the within subject standard error; *p<.05; **p<.01) Microstate D significantly increased GEV between baseline and post-encoding rest. Additionally, there was a significant decrease in the GEV of microstate C.

### 3.4 EEG Spectral Power

In the spectral analysis of the DSOE study, we found two significant clusters. The first cluster consisted included 57 channels and showed a significant increase in power spanning the frequencies 1.17 Hz to 6.84 Hz (57 electrodes, t_sum_ =4.75 × 10^3^, p = 0.013; Figure 5). The second cluster also included all 57 electrodes, but in this case spectral power decreased across the frequency band 8.01 Hz to 12.70 Hz (57 electrodes, t_sum_ =-2.43 × 10^3^, p= 0.050; Figure 5).

**Figure 5:**
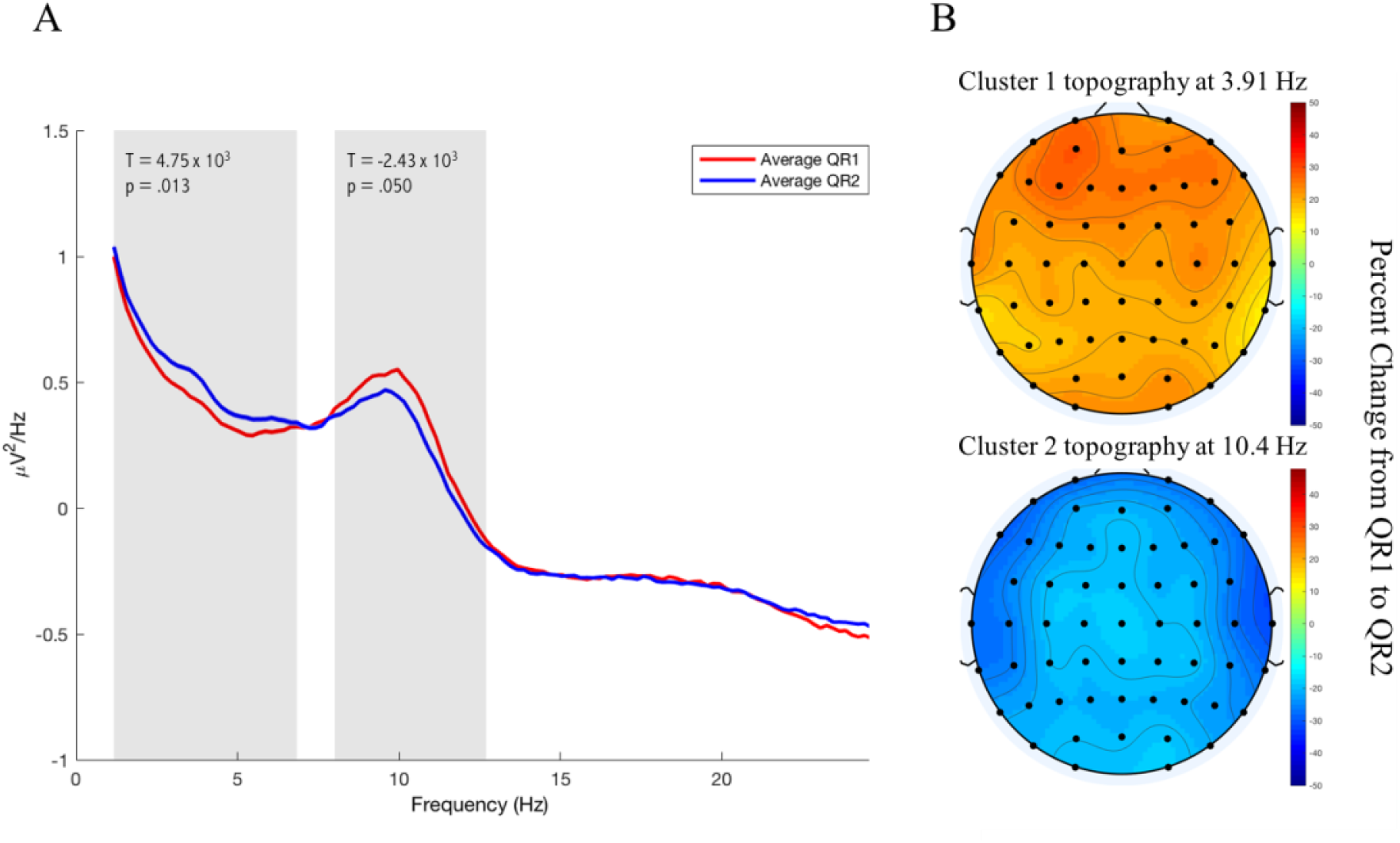
Spectral Power in the DSOE dataset. **A:** The spectral power averaged across all electrodes is plotted in red for QR1 and in blue for QR2. The frequency bands at which the two significant clusters were found are marked by shaded rectangles. **B:** Representative topographies of each cluster are plotted as the percent change in spectral power from QR1 to QR2 at the central value in the range of significant frequencies for each cluster.

### 3.5 Correlations with Memory

Because microstate D showed the most reliable shift in GEV between rest sessions, we focused our memory analysis on this microstate (for an analysis of the relationship between microstate C and memory, see Supplementary Results, Microstate C Correlations with Memory). In the DSOE dataset, microstate D GEV during QR2 significantly correlated with immediate recall for 1 PRES encoded word pairs (r = .35, p = .015; Figure 6, A), as did the increase in microstate D from QR1 to QR2 (r = .32, p = .048; Figure 6, B). When analyzed using a robust linear regression, both effects remained significant (p = .023 and p = .046 respectively. However, neither correlation survived FDR adjustments for multiple comparisons with memory for 1 PRES, 2 PRES, and 4 PRES encoded word-pairs (adjusted p = .069, p = .14 respectively). Significant correlations were not seen with either 2 PRES or 4 PRES encoded word-pair recall (all p’s > .15).

**Figure 6:**
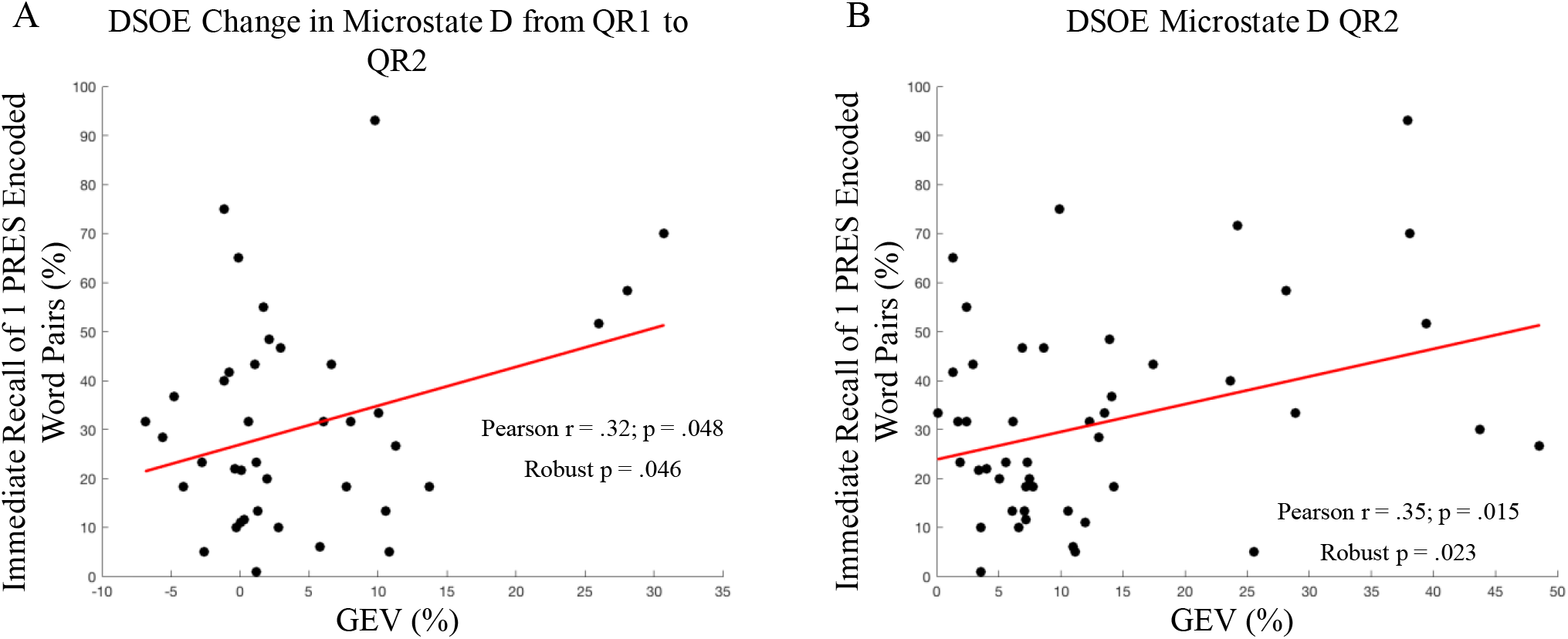
Microstate Correlations with memory. **A:** GEV of Microstate D during post-encoding rest was significantly correlated with memory for 1 PRES encoded word pairs (r = .35). **B:** Microstate D GEV significantly increased from QR1 to QR2; this increase was correlated with memory for 1 PRES encoded word pairs (r = .32).

To further explore the relationship between microstate D and recall performance for 1 PRES encoded word pairs, both effects were analyzed using Meng’s Z-test for comparing correlated coefficients. The relationship between microstate D GEV during QR2 with memory for 1 PRES word-pairs was significantly larger than its correlation with 4 PRES words (p = .022), but not with 2 PRES word-pairs (p = .11). In contrast, the correlation between the increase in microstate D GEV from QR1 to QR2 and 1 PRES word-pairs was not significantly larger than with 2 PRES (p = .17) or 4 PRES (p = .080) word-pairs.

Looking at QR2 spectral power correlations, we found significant clusters that correlated with recall for 1 PRES and 2 PRES word-pairs. The first cluster that correlated with memory for 1 PRES word-pairs spanned all 57 channels and frequencies between 15.23 Hz and 29.88 Hz (57 electrodes, t = 5.15 × 10^3^, p = .035, FDR adjusted p = .052; Figure 7, A). When examining the relationship between spectral power and with recall for 2 PRES word-pairs, we found a second significant cluster which spanned all 57 channels and frequencies between 6.84 Hz and 29.88 Hz (57 electrodes, t = 1.56 × 10^4^, p = .002, FDR adjusted p = .006; Figure 7, B). We found no significant clusters which correlated with memory for 4 PRES word-pairs.

**Figure 7:**
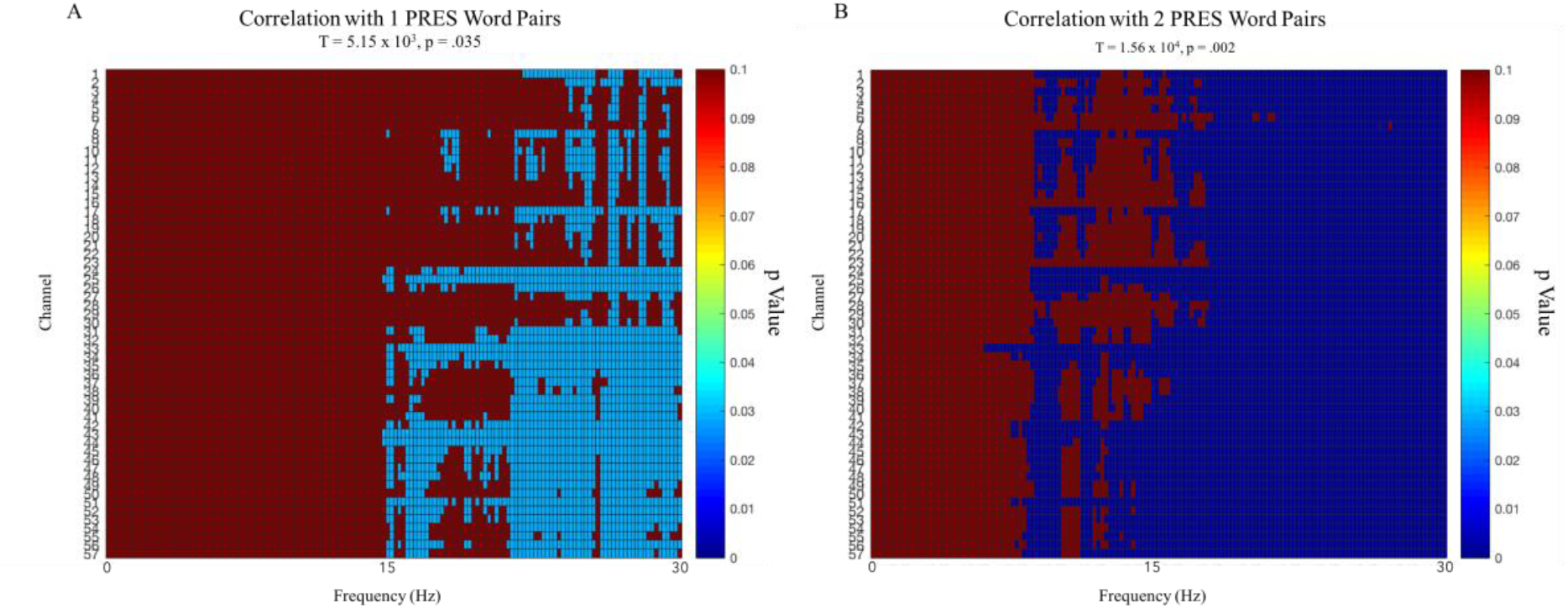
DSOE study. Clusters across channels and frequency significant p values for correlations with 1 PRES and 2 PRES encoded word-pairs. Both clusters ranged across all 57 channels and positively correlated with memory performance.

### 3.6 Alternative Microstate Statistics

Although we chose to focus on microstate GEV in this study, in microstate literature, the statistical measures of duration, occurrence, and coverage are often reported as well. In order to increase the interpretability of our results, we tested our main findings using these other measures in addition to GEV (See Table 1).

**Table 1:** A re-examination of our main findings using the alternate measures of microstate activity: duration, coverage, and occurrence.

| Alternate Measures of Microstate Activity |  |  |  |
| --- | --- | --- | --- |
| Effect | Duration | Coverage | Occurrence |
| Change in microstate D from QR1 to QR2 | Microstate D significantly increased from 63.4ms to 72.8ms ($t(38) = 4.08$, $p = .0002$) | Microstate D significantly increased from 13.1 % to 19.3% ($t(38) = 4.17$, $p = .0002$) | Microstate D significantly increased in appearance from 1.81 times per second to 2.44 times per second ($t(38) = 4.01$, $p = .0003$) |
| Change in microstate C from QR1 to QR2 | Microstate C significantly decreased from 86.6ms to 80.2ms ($t(38) = -2.07$, $p = .045$) | Microstate C showed a trending decrease from 28.7% to 25.2% ($t(38) = -1.95$, $p = .059$) | Microstate C decreased, but not significantly, in appearance from 2.97 times per second to 2.87 times per second ($t(38) = -.76$, $p = .45$) |
| QR2 Microstate D correlation with memory for 1 PRES word-pairs | Microstate D was not significantly related to recall for 1 PRES word pairs ($r = .19$, $p = .19$) | Microstate D was significantly correlated with memory for 1 PRES words ($r = .29$, $p = .04$) | Microstate D was significantly correlated with memory for 1 PRES words ($r = .30$, $p = .04$) |
| Correlation between change in microstate D from QR1 to QR2 and memory for 1 PRESS word-pairs | Change in microstate D did not significantly correlate with recall for 1 PRES word pairs ($r = .16$, $p = .32$) | Change in microstate D was trending towards a significant positive correlation ($r = .29$, $p = .08$) | Change in microstate D did not significantly correlate with recall for 1 PRES word pairs ($r = .14$, $p = .40$) |

Largely, the results using difference statistical measurements seem to either replicate the significance of the effects or trend in the same direction as our findings. Although GEV, duration, coverage, and occurrence are all common microstate measurements, having an effect that remains consistent across all analyses is not guaranteed; for this reason, the relative consistency of our findings across each statistic is an encouraging result.

### 3.7 Stanford Sleepiness Scale

Participants in the DSOE study completed the Stanford Sleepiness Scale (SSS) before beginning the first quiet rest of the protocol. In the DSOE study we found no correlation between the self-reported scores and memory (1 PRES r = -.22, p = .12; 2 PRES r = -.08, p = .57; 4 PRES r = -.16, p = .26). Nor did we find any correlation between SSS scores and microstate GEV in QR1 (Microstate A r = -.16, p = .30; Microstate B r = .08, p = .61; Microstate C r = -.21, p = .18; Microstate D r = .12, p = .45; Microstate E r = .03, p = .87) or in QR2 (Microstate A r = -.07, p = .63; Microstate B r = -.10, p = .50; Microstate C r = .06, p = .67; Microstate D r = -.14, p = .35; Microstate E r = .029, p = .85).

## 4 Discussion

Previous studies have demonstrated that the brain promotes the consolidation of memory during offline periods. Spans of resting wakefulness thus provide a meaningful opportunity to examine how memories are processed by the brain during such periods. In this paper, we discuss a study in which we recorded the EEG activity of participants over the course of rest periods before, between, and after performing a word-pair memory task. In this study, we compared how each subject’s baseline EEG activity changed after learning a series of word-pairs and after being tested on their memory in a cued recall test.

Our investigation of the brain’s electrical activity focused on the use of microstate analysis as a tool to better understand the network dynamics of the memory consolidation process. EEG microstates have been shown to be an effective tool for studying functional networks at a high temporal resolution. Our goal here was to gain a better grasp of the temporal activation of relevant networks, in order to see how this activity changes across the process of learning and consolidation.

### 4.1 Microstate D and C Dynamics Shift During Post-Encoding Rest

The results of our study offer promising insight into the relationship of microstates and memory processing. Here, we confirm the work of previous studies that highlighted the dynamics of microstate D in a post-learning period of resting wakefulness [Murphy et al. 2018]. Indeed, the GEV increase of microstate D during a post-encoding rest was replicated in our study. We further supported these results by using the microstates generated from the DSOE data to analyze the supplementary CES data (see Supplementary Results, Continuous Strength of Encoding Study), where we also found a significant increase in microstate D and decrease in microstate C from QR1 to QR2. When we repeated these analyses using microstates generated from the CES dataset, as well as microstates generated from the combined CES and DSOE data, these results were no longer significant, but showed the same shifting levels of activity across rests.

It is important to note, however, that although our findings reflect the results of *Murphy et al*. 2018, there are several key differences in our study that complicate this interpretation of our findings. First, as we outlined in the Methods section 2.5, the topography of microstate D in *Murphy et al*.’s study had a fronto-central peak, while microstate D in the DSOE data set had an inverse orientation. Although this inverted topography has previously been interpreted as an alternate form microstate D [Michel and Koenig 2018], it is possible that this difference is reflective of two separate microstates that have been grouped together. Second, although *Murphy et al*. included a control condition in which participants went through the experimental protocol without completing the memory task, we had all participants in our study complete the word-pair memory task. Without this control group, we cannot rule out other factors, like arousal, that might have impacted our findings. However, we were able to determine that Stanford Sleepiness Scores were unrelated to either behavior or microstate activity. Finally, the task the subjects performed in *Murphy et al*. 2018 was a virtual maze task that relied on spatial memory as opposed to our word-pair memory task. These memory tasks are not necessarily directly comparable and thus, it is not obvious that our results should replicate the findings in Murphy et al. 2018. This pattern of results across asymmetrical memory tasks could reflect an underlying relationship between microstate D dynamics and memory encoding.

Microstate D has been shown to be associated with the dorsal attention network, a task positive functional network that is more active when performing an attention-focusing exercise [Britz et al., 2010]. Importantly, our study confirms that microstate D is more prevalent in periods of rest after taking part in an encoding task [Murphy et al. 2018]. In contrast to microstate D, microstate C has been repeatedly demonstrated to reflect the activity in the default mode network [Seitzman et al., 2017; Custo et al., 2017; Michel and Koenig 2018]. In the DSOE study, we found a significant decrease in the post-encoding GEV of microstate C as compared to pre-encoding levels. Viewing microstate C activity as indicative of arousal in the DMN, these results suggest that there is a reduction in default mode activity during periods of post-encoding rest. Similarly, Seitzman et al., [2017] illustrated that microstate C is inversely related to task activity, reporting reduced microstate C duration, coverage, and occurrence (GEV not reported)— while completing a serial subtraction task [Seitzman et al., 2017].

The results of our DSOE and supplementary CES analyses provide evidence of an anti-correlated relationship between microstates C and D. Given the growing consensus in the literature of microstate C as indicative of task-negative, DMN activity, and microstate D as emblematic of task-positive network activity [Michel and Koenig, 2018; Seitzman et al., 2017; Custo et al., 2017; Britz et al., 2010; Milz et al., 2016], our results seem to fit with these interpretations of the functional network correlates of microstates C and D. In support of this dichotomous classification, we found a significant negative correlation in the DSOE data between microstate D GEV and microstate C GEV during both the pre-encoding QR1 and the post-encoding QR2. Furthermore, the increase in microstate D from baseline was significantly correlated with the corresponding decrease in microstate C from baseline, suggesting that the dynamics of one microstate are intimately related to those of the other. In concert with existing studies, these results support an inverse relationship between a task-positive microstate D and a task-negative microstate C.

### 4.2 Microstate D is Associated with Memory for 1 PRES Encoded Word-Pairs

Our results show that over the course of quiet rest before and after a period of learning, microstate D fluctuates in ways that correlate significantly with better memory. In the DSOE study, we found that the amount of microstate D activation in QR2 was significantly correlated with memory for 1 PRES encoded words; moreover, the increase in activation of microstate D from QR1 to QR2 correlated with recall for 1 PRES encoded words in the DSOE study. This suggests that activation of microstate D, and putatively the dorsal attention network, may be related to the consolidation of weakly encoded information. This result seemingly contrasts with the finding in Murphy et al. [2018], in which microstate D was negatively associated with improvement in a post-nap maze task. However, in Murphy et al.’s study, for subjects performing a virtual maze task, as opposed to a word-pair memory task, microstate D was only negatively correlated with *post-nap* memory performance [Murphy et al., 2018]. Our current study examined the immediate recall of recently learned information after a short period of quiet rest, which may differ decidedly from the effect sleep-based memory consolidation has on recall performance. In this case, microstate D may play a role in the immediate consolidation of weakly encoded material that diverges from its impact on later sleep-based memory consolidation.

The correlations between microstate D and memory, may aid in the understanding of how encoding strength modulates the consolidation of memories. Our results from the DSOE data suggest that increased microstate D GEV, and the magnitude of the GEV increase, are associated with better memory for 1 PRES encoded words. These findings are in line with the plethora of research in the sleep and memory literature that indicate the prioritization of weakly encoded memories during sleep-based consolidation [Drosopoulos et al., 2007; Schapiro et al., 2017; Djonlagic et al., 2009; Denis et al., 2019; Denis et al., 2020].

Although the changes in microstate D and C were replicated in both the DSOE and the supplementary CES datasets, our preliminary findings of an association between cued recall and both post-encoding microstate D and the overall change in microstate D from baseline merit further exploration. Additionally, the suggestion that encoding strength could play a role in the prioritization, during quiet rest, of memories for subsequent consolidation presents an interesting starting point for future studies.

### 4.3 Spectral Analysis Leads to Mixed Results

The spectral analysis of the DSOE data showed a significant increase in power in frequencies spanning the delta and theta bands in the DSOE study while significantly *decreasing* across frequencies in the alpha band. Despite this overall decrease in power, we also found that greater power in alpha frequencies correlated significantly with memory recall (see Supplementary Results, Spectral Power Correlations with Memory). However, Brokaw et al. [2016] found the inverse of this relationship between alpha oscillations and memory retention, reporting increased verbal memory was associated with lower alpha power in post-learning rest [Brokaw et al., 2016]. Interestingly, Murphy et al., [2018] also found a significant increase in alpha power during a rest period after training on a spatial memory task [Murphy et al., 2018]. The contradictory results of our studies are seemingly reflective of a greater discrepancy throughout the literature on this subject.

## 5 Limitations

A few key limitations exist in our study and analysis that are important to note in order to interpret these data. First, our study design lacked a non-rest control group. Without this control, these data cannot definitively show that quiet rest benefits memory, only that, within a quiet rest, the specified EEG patterns predict recall performance. Moreover, we also did not control for arousal effects on EEG activity or memory recall by adding a control-group that did not perform the memory task. While we did not find any correlations between self-reported arousal and either microstate dynamics or memory performance (see 3.7 Stanford Sleepiness Scale), without this control group, we cannot sufficiently rule out any influence of arousal on neural activity or recall accuracy. Finally, while we did control for the length of words across encoding groups, we did not control for the average number of syllables across word-pair groups. Controlling the average number of syllables within an encoding group could help ensure parity in the difficulty of the memorization task across groups of word-pairs.

This analysis is strengthened by the use of a separate supplementary dataset, in which we were largely able to replicate results across both studies. One drawback, however, is that our manipulation of encoding strength was not necessarily consistent across the two experiments—in fact, L PRES encoded word-pairs in the CES study seemed to be equivalent to 1 PRES encoded pairs from the DSOE study. With this in mind, it is difficult to directly compare the behavioral results from the DSOE and CES studies.

In order to fully understand the relationship between the EEG dynamics of a post-encoding quiet rest and memory, future studies should take these concerns into account.

## 6 Conclusion

The results of this study demonstrate the value of performing EEG microstate analyses in studies of neurological activity during periods of memory consolidation. Given the interpretation of microstates as instances of *quantized cognition*, they provide a unique insight into cognitive processes at a high temporal resolution. In this study, we were able to consistently replicate previous important findings on the post-learning increase in the GEV of microstate D. Our data help to reinforce the current view that Microstate D reflects the activity of a task-positive network related to the dorsal attention system and that Microstate C is representative of activation of a task-negative system akin to the DMN. Here we hope that we have raised intriguing questions for the field of memory consolidation and that we have further provided a direction for future research into the relationship between microstate D and memory consolidation and enhancement.

## Supporting information

Supplementary Results

## Acknowledgments

This study was supported by N.I.H. grant MH 048832. We thank Alex Morgan, Elaine Parr, Verda Bursal, and the staff of the Clinical Research Center at Beth Israel Deaconess Medical Center for their technical support.

