## Supplementary Results for "Using EEG Microstates to Examine Post-Encoding Quiet Rest and Subsequent Word-Pair Memory"

### Continuous Strength of Encoding Study

#### Methods

##### Study 2—Continuous Strength of Encoding

Participants:

Twenty Subjects (14 female, mean age = 24, SD = 4.34 years) took part in this protocol; as in Study 1, one subject was excluded for having been unable to correctly remember any word pairs from the learning task. Additionally, due to the addition of a baseline quiet rest partway through data collection, 5 subjects did not have a quiet rest 1. Participants in this study underwent the same screening procedure as in Study 1. As in that study, they were instructed to maintain a regular sleep schedule for 3 nights leading up to the study and to refrain from consuming any caffeine on the morning of the experiment. Recruitment was through online postings at local universities advertising a learning and memory study. Participants were compensated for their time. The procedures were approved by the institutional review board of Beth Israel Deaconess Medical Center.

**Procedure:**

Participants began the protocol at 9:00pm, providing informed consent and then completing two surveys: the first evaluated their quality of sleep over the past three nights, while the second asked about their sleep over the past month (Pittsburgh Sleep Quality Index (PSQI)).

After being fitted with an EEG cap, they were given 5 minutes of eyes-closed quiet rest (QR1). The subjects then began the encoding session, in which they were shown each word pair for a different amount of time in order to produce unique encoding strengths for each word pair. Immediately after the study phase, subjects were given a second 5-minute period of eyes-closed rest (QR2), followed by initial testing of their memory for the word pairs (Supplementary Figure 1). As with Study 1, the protocol continued after this rest. Data collected beyond this point are the subject of a separate analysis.

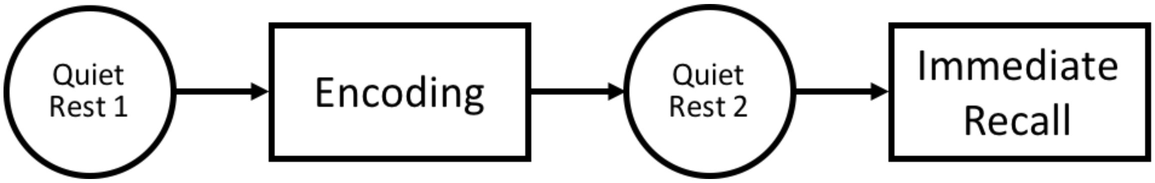


**Supplementary Figure 1:** Continuous Strength of Encoding (CES) study protocol. Subjects began with a 5-minute baseline rest before being presented with 90 word-pairs for varying periods of time, ranging from 50ms to 4,500ms. After a post-encoding 5-minute rest, subjects were tested on their memory for the word-pairs. Analyses of the tasks performed after this protocol are analyzed in a separate paper.

**Encoding and Recall Task:**

This study dovetails with the Discrete Strength of Encoding study by looking at encoding strength as a continuous rather than discrete variable. Participants were asked to complete a word-pair memory task, this time, with 90 word-pairs. However, each word pair was presented on screen for a different length of time ranging from 50ms to 4,500ms in steps of 50 milliseconds. Although the smallest presentation time was 50ms, analysis was only performed on presentation times of 800ms and longer as participants largely failed to recall word pairs presented for shorter times. To compare the results between studies, the word-pair durations were divided into short "S PRES" (800-2,000ms), intermediate "I PRES" (2050-3,250ms), and long "L PRES" (3,300-4,500ms) subsets of 25 word pairs each.

#### Results

##### Behavior

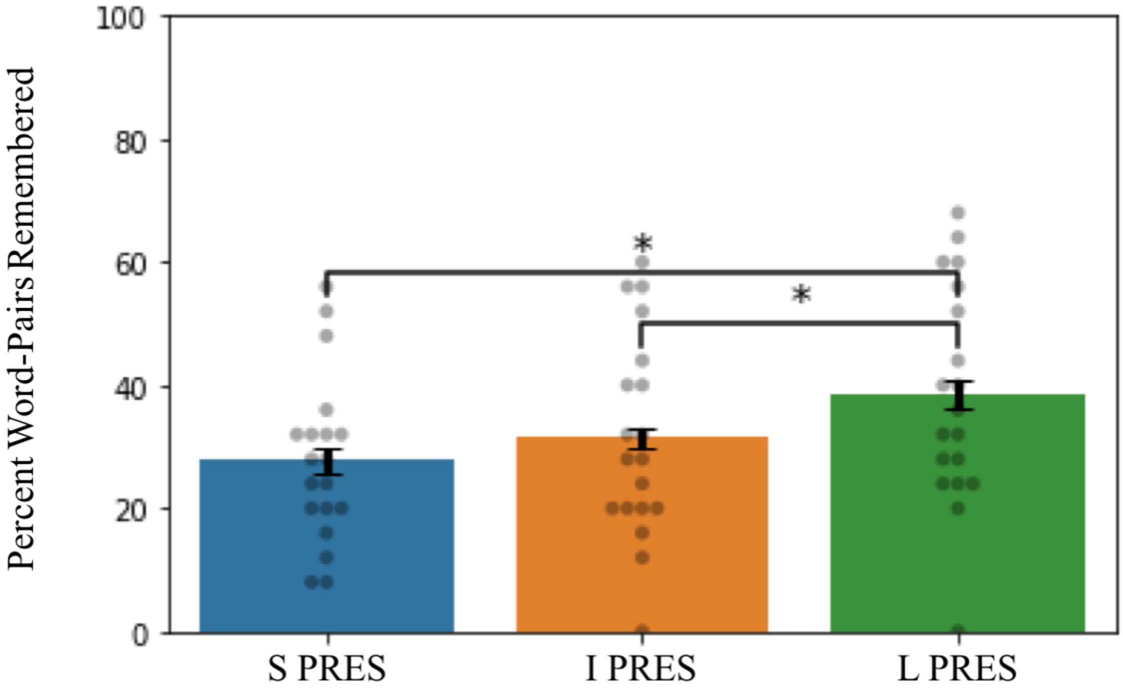


**Supplementary Figure 2:** Memory performance in the CES study (error bars display the within subject standard error; dots represent individual subject values; *p < .05).

The same pattern of results found in the DSOE study was obtained in the CES study, albeit with poorer performance overall. L PRES encoded words were remembered most effectively at 39.4%, while I PRES and S PRES encoded word-pair performance was 32.6% and 28.0% respectively. L PRES encoded words were recalled significantly better than either S PRES ($t(18) = 2.9$, $p = .010$, FDR adjusted $p = .038$) or I PRES ($t(18) = 2.2$, $p = .039$, FDR adjusted $p = .053$) encoded words, though the difference between L PRES and I PRES encoded words was only trending towards significance after FDR adjustment (Supplementary Figure 2). Although, L PRES encoded word-pairs were remembered significantly better than S PRES or I PRES encoded words, overall memory for L PRES encoded word in the CES study was on par with performance for 1 PRES encoded words in the DSOE study—in fact, using a two-sample t-test, the two groups were statistically indistinguishable (t(70) = 1.30, p = .20).

**Microstate fit**

The microstate fit was lower for the CES study, presumably because the microstate maps were generated from the DSOE study. On average, the five topographies explained 51.7% of the global variance of CES subjects in QR1 and 59.6% in QR2.

**Microstate Dynamics Across Periods of Rest**

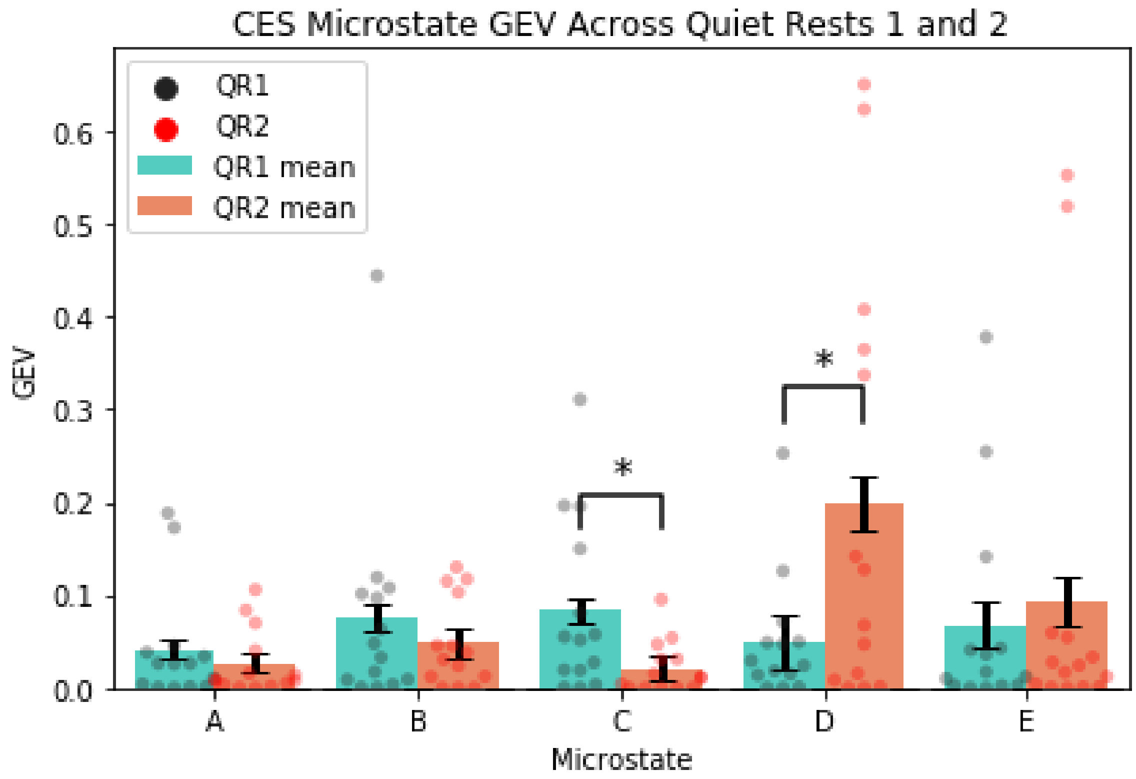

**Supplementary Figure 3:** Measuring GEV across rest periods (error bars display the within subject standard error; colored dots represent individual subject values across conditions; *p<.05; **p<.01) Microstate D significantly increased GEV between baseline and post-encoding rest. Additionally, there was a significant decrease in the GEV of microstate C. These results replicate the microstate C and D findings in the DSOE dataset.

We were able to replicate the DSOE dataset findings in the CES dataset. When we fit the DSOE-based microstates to the CES EEG records, we replicated both the microstate D and C findings seen in the DSOE dataset (Supplementary Figure 3). We found a significant increase in the GEV of microstate D from 5.0% in QR1 to a post encoding rest 19.9% in QR2 ($t(13) = 2.6$, $p = .023$), which was present in 79% of subjects. Furthermore, the decrease in microstate C from 8.3% to 2.1% ($t(13) = -2.5$, $p = .026$) was also significant and was present in 93% of participants. Both results reflect the effects found in the DSOE data. However, after a post-hoc statistical power analysis, we determined that that the CES dataset was underpowered to achieve 80% confidence in our replication of the change in microstate C (power = .64) and microstate D (power = .68) from QR1 to QR2.

Additionally, only the correlation between microstates C and D GEV during QR2 ($r = -.34$, $p = .16$, robust regression $p = .34$) was consistent with the DSOE study. The relationship between microstates C and D during QR1 ($r = .08$, $p = .80$, robust regression $p = .97$) and the shift in GEV from QR1 to QR2 ($r = .02$, $p = .94$, robust regression $p = .51$) contrasted with the results from the DSOE dataset.

**EEG Spectral Power**

In the CES data, we found one significant cluster showing increased power of 56 channels (all but Fp1) in the range of 9.18 Hz to 29.88 Hz ($t_{sum}$ = 6.25 x $10^3$, p = .03; Supplementary Figure 4).

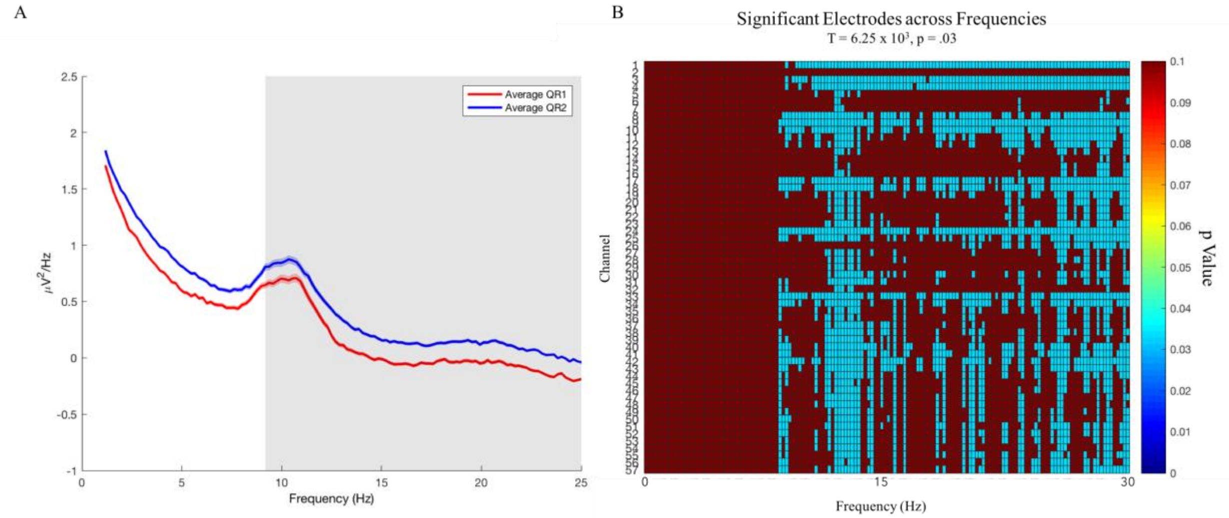


**Supplementary Figure 4:** Spectral power in the CES dataset. **A:** The spectral power averaged across all electrodes is plotted in red for QR1 and in blue for QR2. The frequency bands at which the significant cluster was found is marked by a shaded rectangle. **B:** Cluster across channels and frequency significant p values. This cluster ranged across all channels except for Fp1 and spanned the frequencies 9.18 Hz to 29.88 Hz.

#### Correlations with Memory

In the CES dataset, we found no significant correlations between QR2 microstate D GEV with S PRES (r = .28, p = .25), I PRES (r = .13, p = .60), L PRES (r = .32, p = .18). We did, however, find that the increase in microstate D from baseline significantly correlated with recall for L PRES encoded words (r = .59, p = .028, robust p = .013, FDR adjusted p = .039; Supplementary Figure 5). The correlations with S PRES (r = .39, p = .17, robust p = .23) and I PRES (r = .22, p = .44, robust p = .459) were not significant.

To further test these results, we used Meng's Z-test for correlated coefficients to examine whether the correlation between the increase in microstate D GEV and L PRESS words was significantly greater than its correlation with either S PRES or I PRES word-pairs. We found that the correlation between the increase in microstate D GEV and L PRES word-pairs was not significantly larger than its correlation with S PRES word-pairs (p = .22), and was only trending towards significantly larger than its correlation with I PRES word-pairs (p = .06).

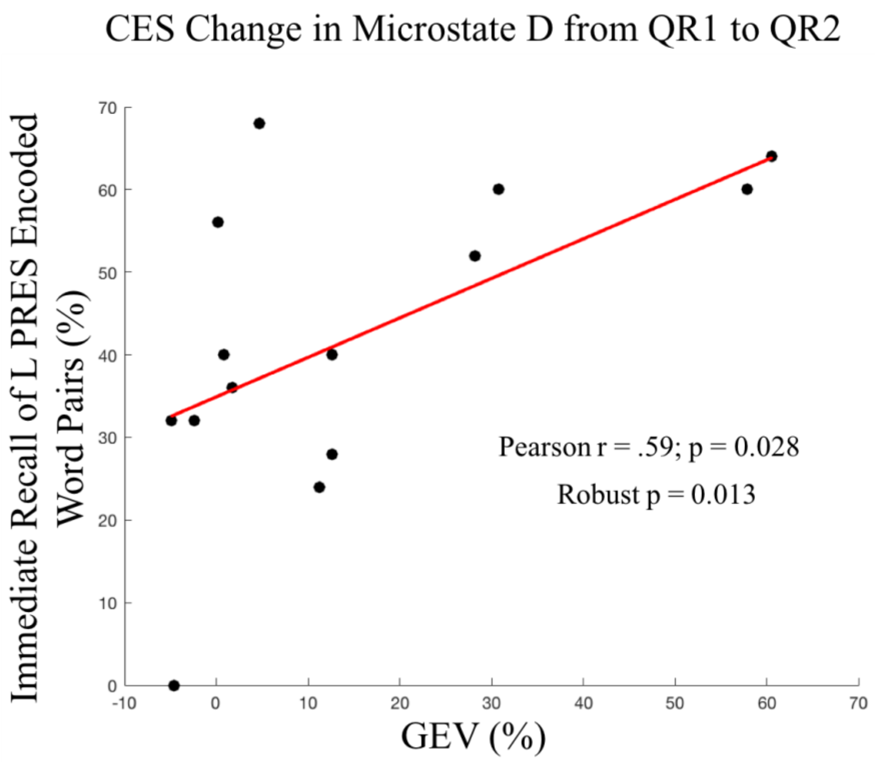


**Supplementary Figure 5:** In the CES study, the increase in microstate D GEV from baseline was positively correlated with recall for L PRES encoded words (r = .59).

**<u>Alternate Microstate Topographies:</u>**

<u>CES Topographies</u>

After analyzing the CES data using the microstate topographies generated from the DSOE study, we sought to replicate our findings from microstates that were generated using the CES data. After using the same procedure detailed in Methods section 2.5, we were able to generate a new set of five microstates (Supplementary Figure 6). We opted to use 5 topographies in order maintain consistency with the DSOE study. While these new topographies contained all four canonical microstates, they did not directly match the 5 microstates generated from the DSOE study, specifically, microstate E is no longer present.

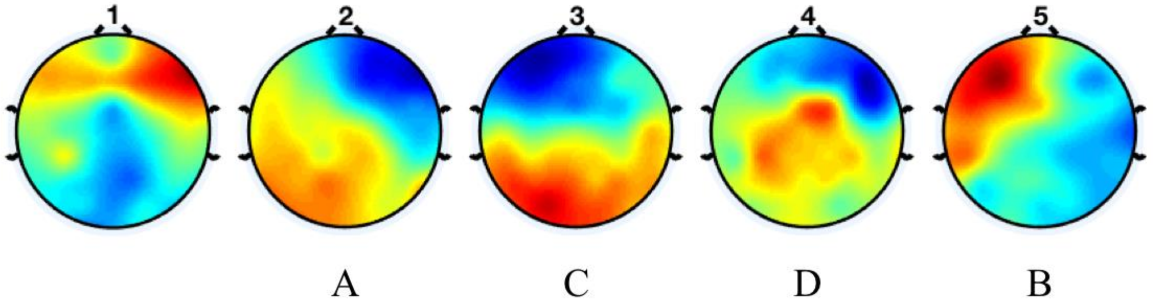


**Supplementary Figure 6:** Microstate topographies generated from the CES dataset. All four canonical microstates were present, however microstate E from the DSOE dataset was no longer present.

Surprisingly, the fit for these new topographies was marginally lower (QR1 = 53.07 % GEV and QR2 = 58.19 GEV) than the fit for the DSOE topographies. This contradictory finding seems to suggest that our small sample size in the CES dataset is not sufficient to produce a regularized set of topographies that better fits the available data. Additionally, after computing the microstate statistics for QR1 and QR2, we found no significant differences across rests in the GEV of any microstates (A: t(13) = 1.50, p = .15; B: t(13) = -

1.00, p = .33; C: t(13) = -1.63, p = .13; D: t(13) = .83, p = .42; Microstate 1: t(13) = .98, p = .34; Supplementary Figure 7). Although these relationships were not significant, the pattern of results did still show an increase in microstate D (from 1.58% to 5.90%) and a decrease in microstate C (from 20.65% to 12.04%) across rest periods.

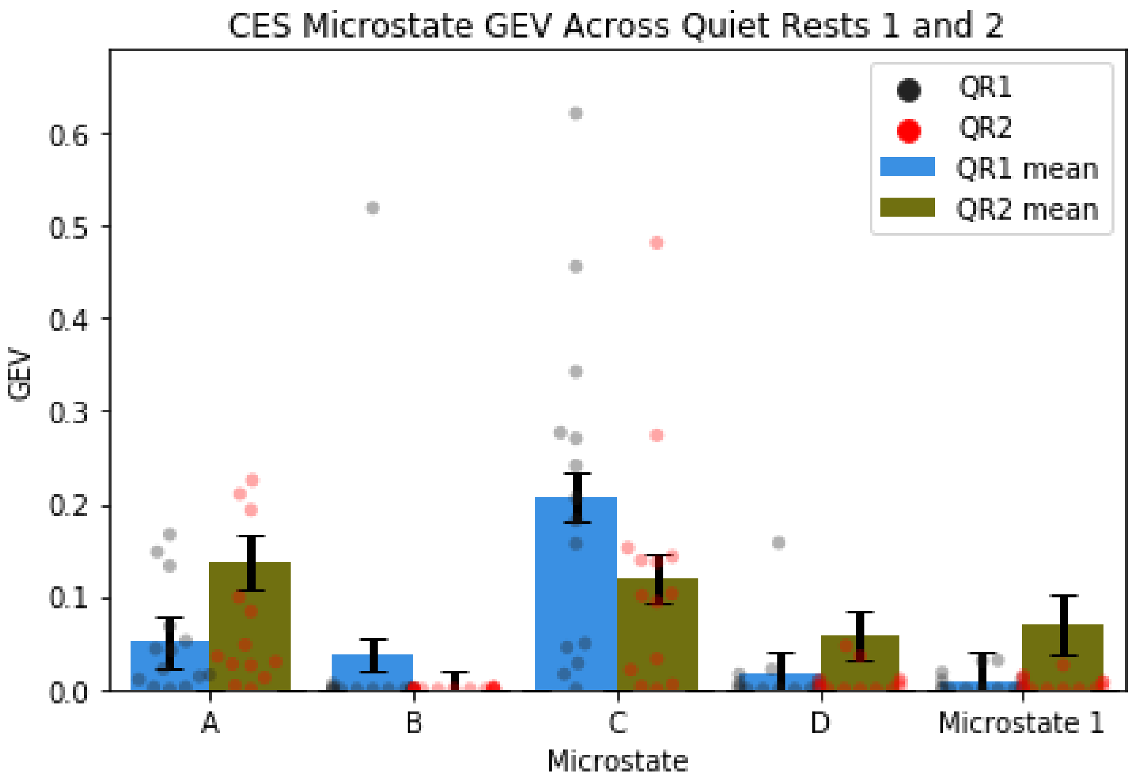


**Supplementary Figure 7:** Measuring GEV across rest periods using microstates generated from the CES data (error bars display the within subject standard error; colored dots represent individual subject values across conditions). Although there were no significant changes across QR1 and QR2, there is still an increase in microstate D GEV and a decrease in microstate C GEV, which is reminiscent of the effect we found using the DSOE microstates.

Finally, we looked at the relationship between the CES microstate D and memory for word-pairs across encoding groups. We found a significant positive correlation between QR2 microstate D for S PRES word-pairs (r = .48 p = .035, FDR corrected p = .10) but not for

I PRES (r = .28 p = .25, FDR corrected p = .25) or L PRES word-pairs (r = .16 p = .51, FDR corrected p = .51). We also found a significant correlation between the increase in microstate D from QR1 to QR2 and S PRES word-pairs (r = .57 p = .035, FDR corrected p = .11) but not for I PRES (r = .26 p = .35, FDR corrected p = .53) or L PRES word-pairs (r = .15 p = .60, FDR corrected p = .60). Although these relationships did not survive FDR correction, they do reflect the relationship that we found in the DSOE dataset between QR2 microstate D and the increase in microstate D across rests with memory for 1 PRES words.

Combined CES and DSOE Topographies

In addition to generating microstates from the CES data, we also generated a set of microstate topographies using the normalized combination of the CES and DSOE datasets. In order to combine the CES and DSOE datasets without signal flattening (see Signal Flattening), additional channels were interpolated in the re-referenced CES data and 13 records (5 QR1 and 8 QR2) were removed. This left us with 9 QR1 records and 11 QR2 records to analyze.

The microstates created from the combined data (henceforth, combined microstates) had a higher fit for the CES data than either the microstates generated from the DSOE or CES data separately (QR1 = 71.55% GEV and QR2 = 70.08% GEV). All four canonical microstate topographies were present, though microstate D was notably skewed on a slight rightward angle (distinct from the orientation of microstate A) (Supplementary Figure 8). The fifth and final microstate did not match any previously seen microstate topographies.

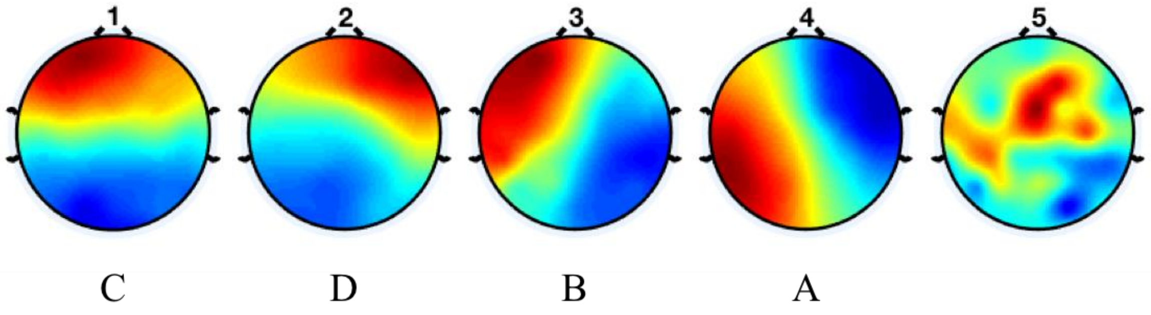


**Supplementary Figure 8:** Microstate topographies generated from the combined CES and DSOE dataset. In these combined microstates, the canonical topographies of A, B, and C are all clearly present; microstate D however, seems to be tilted in a manner that is atypical for its usual orientation.

We found no significant differences in the GEV of any microstates between QR1 and QR2 (A: $t(6) = -.45$, $p = .67$; B: $t(6) = 1.04$, $p = .34$; C: $t(6) = -.60$, $p = .57$; D: $t(6) = 1.44$, $p = .20$; Microstate 5: $t(6) = -.97$, $p = .37$; Supplementary Figure 9). We did, however, see a decrease in microstate C (from 23.51% GEV to 17.50% GEV) and an increase in microstate D (5.09% GEV to 16.16% GEV) from QR1 to QR2.

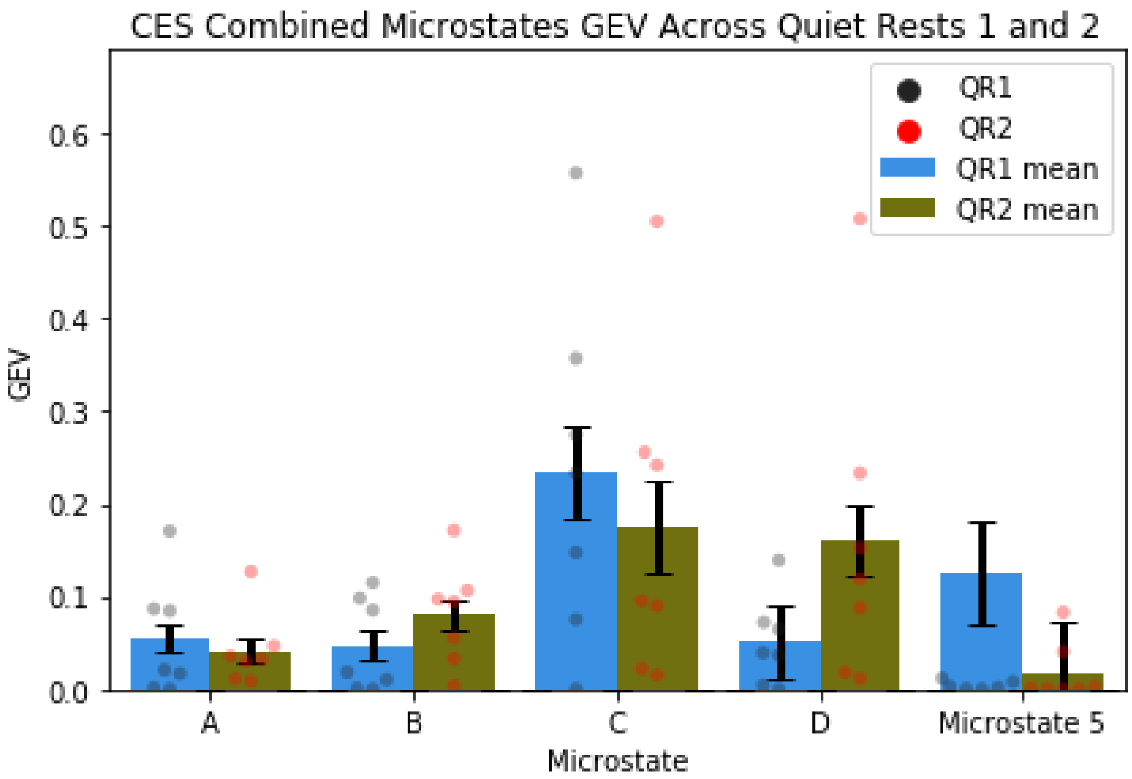


**Supplementary Figure 9:** Measuring GEV across rest periods using microstates generated from the combined CES and DSOE data (error bars display the within subject standard error; colored dots represent individual subject values across conditions). Although there were no significant changes across QR1 and QR2, we still find an increase in microstate D GEV and a decrease in microstate C GEV, which is reflects our findings in the DSOE data and is consistent across the CES data regardless of microstate topographies.

Finally, we found a significant positive correlation between microstate D GEV during QR2 and memory for S PRES word-pairs (r = .64, p = .034, FDR corrected p = .10), but not for I PRES (r = .48, p = .14, FDR corrected p = .18) or L PRES (r = .44, p = .18, FDR corrected p = .18) word-pairs. Although this relationship did not survive FDR correction, it did replicate our findings in the DSOE dataset and in the CES dataset using CES microstate topographies. In all three cases, we found a positive correlation between QR2 microstate D and memory for word-pairs in the weakest encoding group. We found no correlations between the change in microstate D from QR1 to QR2 and memory for word-pairs in any encoding group (S PRES: r = .62, p = .13, FDR corrected p = .40; I PRES: r = .13, p = .78, FDR corrected p = .81; L PRES: r = .11, p = .81, FDR corrected p = .81).

#### **Discussion**

In the CES study, the increase in microstate D from QR1 to QR2 was correlated with memory for L PRES encoded words.  It is worth noting that overall memory for word-pairs was much worse in the CES study that in the DSOE study; furthermore, memory for L PRES encoded words in the CES study (39.4%) was not significantly different from memory for 1 PRES encoded words in the DSOE study (31.7%). To that end, L PRES encoding in the CES study set may be akin to 1 PRES encoding in the DSOE study, in which case we would expect to see such a correlation with the increase in microstate D. One factor for consideration is that the smaller sample size of the CES dataset, as compared to the DSOE dataset (14 vs. 52 subjects), increases the likelihood of spurious correlations, which could explain why the results were not consistent across the two studies.

### Additional Results for Both DSOE and CES Studies

**Channel Interpolation and Component rejection:**

In the DSOE study, the average number of channels interpolated was 7.48 (SD 8.86) and the average number of rejected components was 2.13 (SD 0.94). In comparison, the supplementary CES study the average number of channels interpolated was 4.14 (SD 7.14) and the average number of rejected components was 2.0 (SD 1.0). The DSOE data, however, was also reexamined and additional channels were interpolated after the data were re-referenced to the average across electrodes. This was in order to combat the signal flattening that we observed in the generated microstates (see Signal Flattening).

**Signal Flattening:**

Some EEG records were removed from the analysis due to a flattening of the averaged reference signal. When we examined these records, they would produce microstates with a global topography of 0mV.

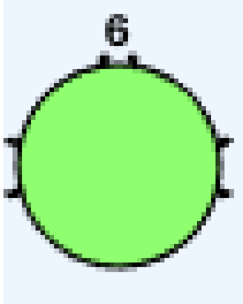


Example from subject run that was excluded.

**Microstate Fit:**

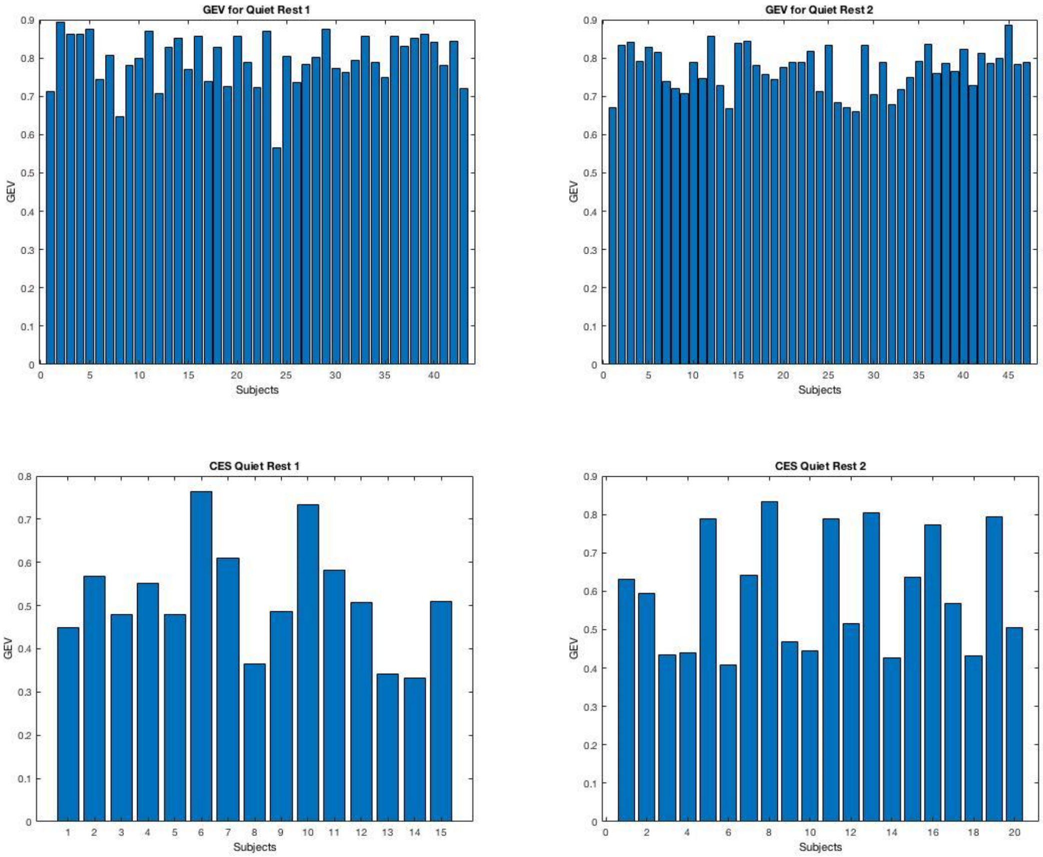


**Supplementary Figure 10:** GEV across individual subjects for quiet rest sessions in the DSOE study (top row) and the CES study (bottom row).

**Quiet Rest 3 Results:**

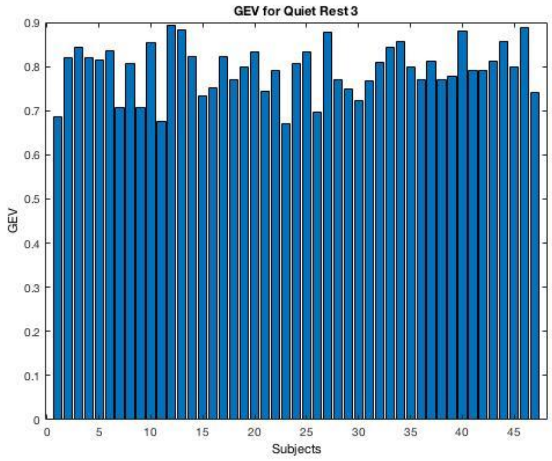


**Supplementary Figure 11:** In QR3, the 5 template microstates explained 79.40% of the global variance.

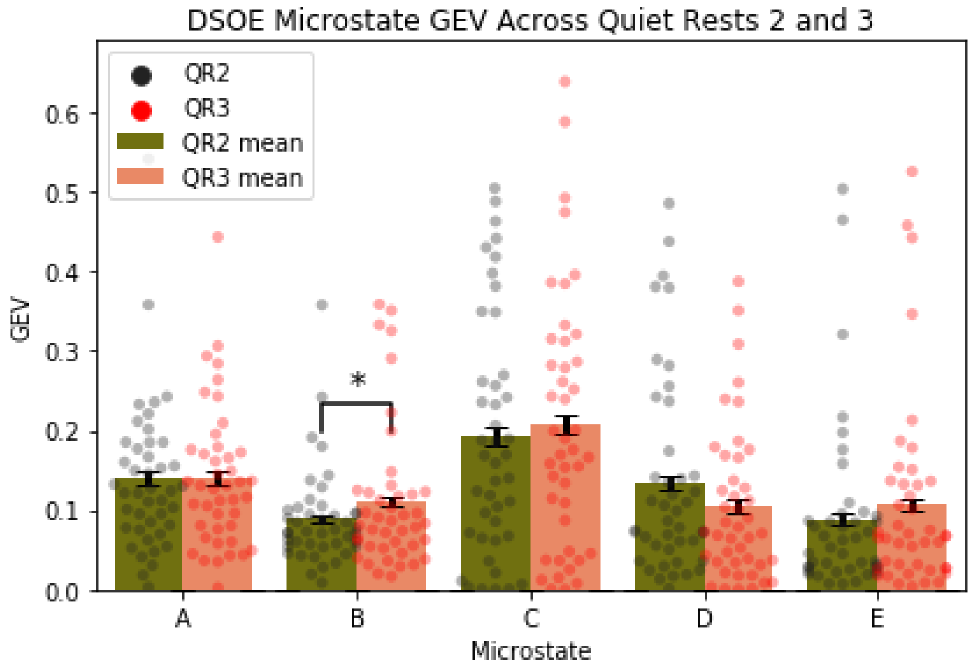

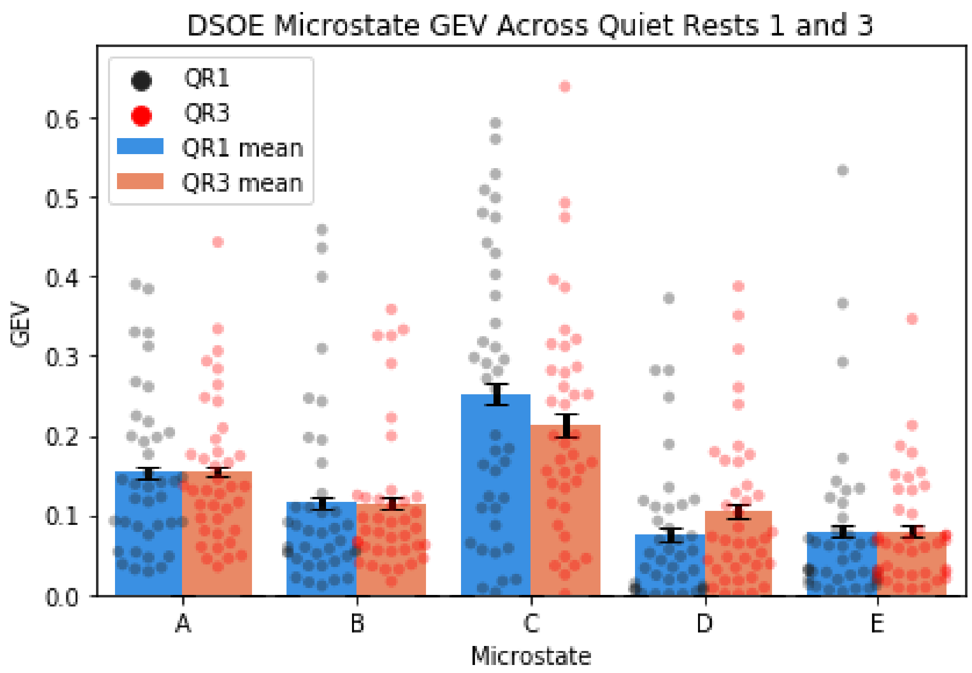


**Supplementary Figure 12: Top:** Microstate B significantly increased in GEV from quiet rest 2 to quiet rest 3 (p = .0481). **Bottom:** There was no statistical difference in the amount of any microstate in baseline rest and quiet rest 3(error bars display the within subject standard error; colored dots represent individual subject values across conditions; *p<.05).

**Relationship between Microstate C and D in the DSOE Study:**

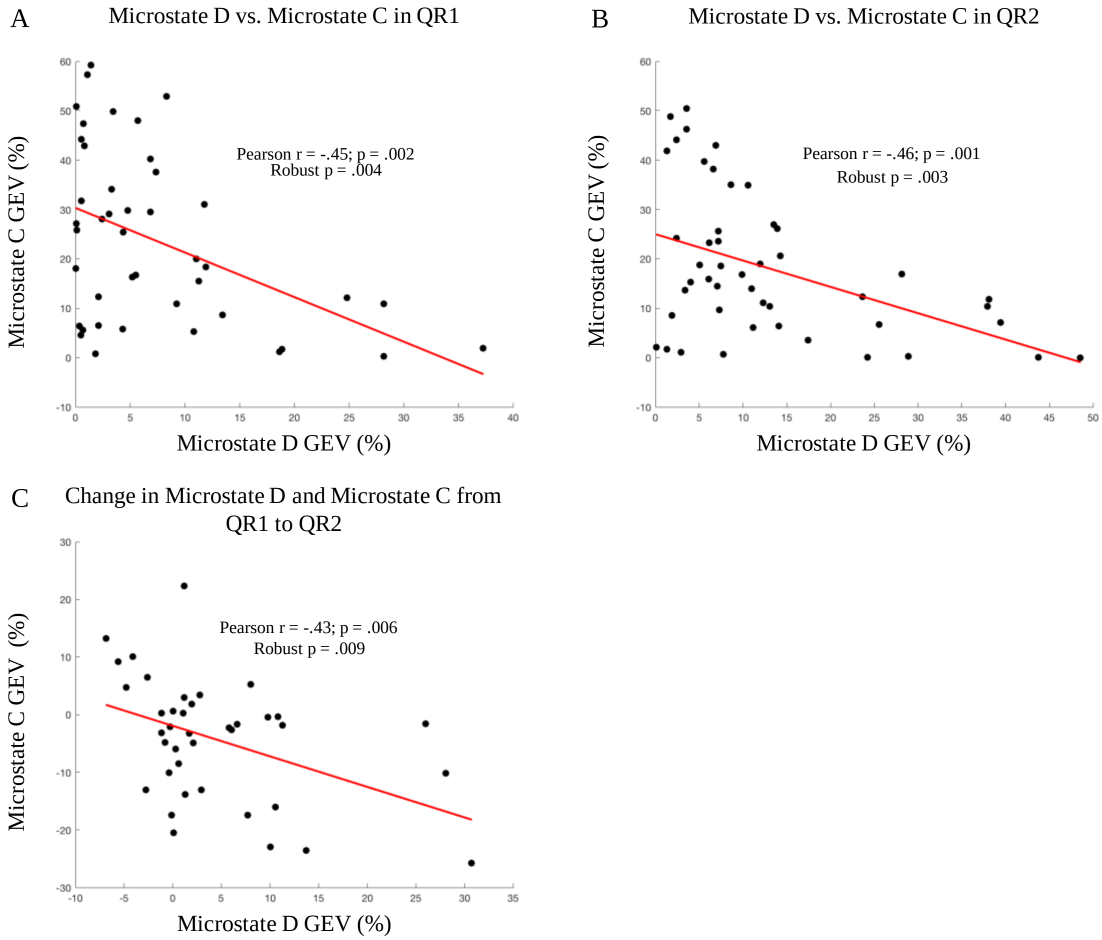


**Supplementary Figure 13: A:** Microstate C GEV was negatively correlated with Microstate D GEV during QR1. **B:** GEV of Microstate C and D was also negatively correlated during QR2. **C:** Changes in Microstate C and D from QR1 to QR2 were also significantly negatively correlated.

In order to fully analyze the relationship between microstates C and D, we next examined whether the ratio of microstate D to microstate C GEV in QR2 was correlated with memory performance across encoding categories. We found no significant correlations between the ratio of microstates C and D with any encoding category (1 PRES r = -.030, p =

.84; 2 PRES r = -.018, p = .91; 4 PRES r = .060, p = .69); nor did we find any relationship between memory and the change in the ratio of microstate D to C from QR1 to QR2 (1 PRES r = -.037, p = .83; 2 PRES r = -.014, p = .93; 4 PRES r = .068, p = .68).

**Microstate C Correlations with Memory:**

In the DSOE study we also observed a significant decrease in the GEV of microstate C from QR1 to QR2. However, post-hoc tests revealed no significant relationship between microstate C and recall performance in either the DSOE dataset. Microstate C GEV during QR2 was not significantly correlated with memory for weak (r = -.12, p = .44, robust regression p = .53), intermediate (r = .04, p = .81, robust regression p = .64), or strong (r = -.18, p = .218, robust regression p = .89) encoding pairs. The decrease in microstate C from QR1 to QR2 was also not significantly correlated with weak (r = -.22, p = .18, robust regression p = .17), intermediate (r = -.25, p = .13, robust regression p = .22), or strongly encoded word-pair memory (r = -.33, p = .041, robust regression p = .65).

Similarly, in the CES study, microstate C GEV during QR2 was not significantly correlated with memory for intermediate (r = -.1, p = .69, robust p = .66) or strong (r = -.30, p = .22, robust p = .28) encoding pairs. Finally, the decrease in microstate C from QR1 to QR2 was not significantly correlated with intermediate (r = .13, p = .65, robust p = .56) or strongly encoded word-pair memory (r = .21, p = .48, robust regression p = .49). However, after using a robust regression, we found that both microstate C GEV during QR2 (r = -.10, p = .68, robust p = .001, FDR adjusted p = .003; Supplementary Figure 14, A) and the decrease in microstate C GEV (r = -.21, p = .47, robust p = .027, FDR adjusted p = .081; Supplementary Figure 14, B) were correlated with memory for weakly encoded words. It should be noted that only the relationship between microstate C GEV in QR2 and recall of weakly encoded word-pairs remained significant after performing an FDR adjustment.

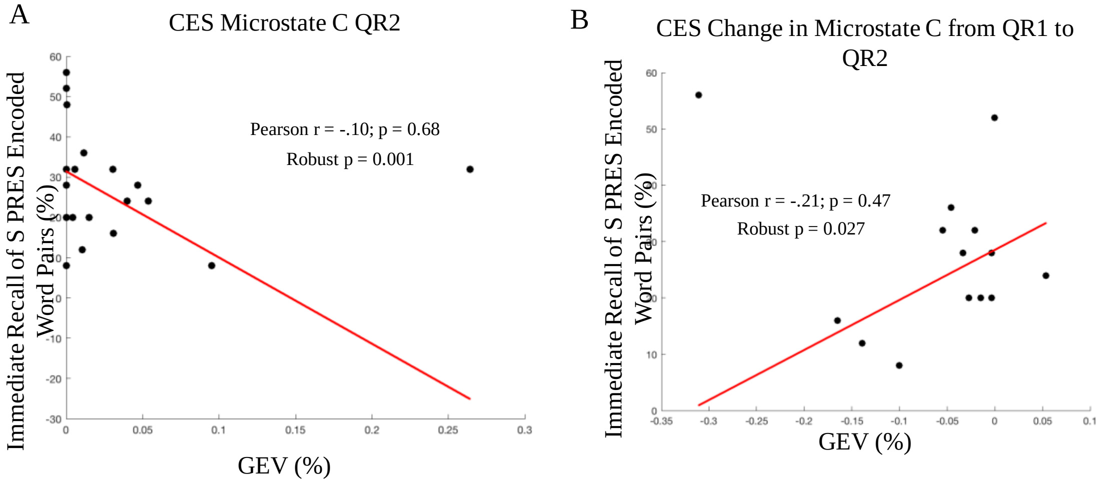


**Supplementary Figure 14: A:** After a robust regression, microstate C GEV during was negatively correlated with memory for S PRES encoded words in the CES study. **B:** The decrease in microstate C from QR1 to QR2 in the CES study was positively related to memory for S PRES encoded words.

#### Spectral Power Correlations with Memory:

There were no significant clusters associated with memory performance in the CES dataset.

#### CES Stanford Sleepiness Scale:

Paralleling the DSOE study, participants in the CES study also completed the Stanford Sleepiness Scale (SSS) before beginning the first quiet rest of the protocol. In the CES dataset, we also did not find correlation between the SSS scores and memory (S PRES r = .09, p = .72; I PRES r = .18, p = .47; L PRES r = .12, p = .63). Similarly, we found no relationship between SSS values and microstate GEV in QR1 (Microstate A r =-.48, p = .08;

Microstate B r = .28, p = .34; Microstate C r =.27, p = .36; Microstate D r = .29, p = .31; Microstate E r = -.09, p = .74) or QR2 (Microstate A r = .30, p = .22; Microstate B r =-.20, p = .42; Microstate C r = .11, p = .66; Microstate D r = .08, p = .75; Microstate E r = -.10, p = .67).
